# Spatial density and habitat associations of Atlantic Cod on the Northeastern US Continental Shelf

**DOI:** 10.1101/2025.01.08.631947

**Authors:** Katie Lankowicz, Graham D. Sherwood, Jonathan H. Grabowski, Lisa Kerr

**Affiliations:** Gulf of Maine Research Institute Portland, Maine; Northeastern University Marine Science Center Nahant, MA; University of Maine School of Marine Sciences Orono, ME

**Keywords:** Atlantic Cod, vector autoregressive spatiotemporal model, spatial density, habitat

## Abstract

The spatial distribution of the Atlantic cod (*Gadus morhua*) stock is shaped in part by several habitat and oceanographic variables. In this study, Vector Autoregressive Spatio-Temporal (VAST) models were used to combine data from multiple survey programs to hindcast seasonal spatial densities of three size classes of cod within the Northeast US Continental Shelf from 1982 to 2021. Bottom habitat characteristics, bottom water temperature, depth, and basin-averaged climate indices were included as density covariates. Depth, bottom temperature, and gravel sediments were strongly associated with spatial density. The relative abundance of all size classes generally decreased throughout the time series. Model outputs highlighted patches with persistently high spatial density despite range losses and declining abundance. This aligns with the basin model, a spatial dynamic frequently reported in collapsed fish stocks. The availability of habitat with suitable depth and temperature will likely be reduced under current projections of bottom water temperature, further endangering the recovery of the stock. Improving our understanding of cod habitat preferences and variation in spatial density will be important for future management efforts.

## 1 Introduction

Atlantic cod (*Gadus morhua*) are an ecologically important part of New Englands groundfish complex, and an economically and culturally important part of New Englands groundfish fishing industry. Once a mainstay of the commercial fishery, the reduced quotas of cod now limit the exploitation of other species in the groundfish complex. Atlantic cod stock assessments and management efforts in the Northeast US continental shelf (NEUS) region are informed by a suite of fishery-independent surveys orchestrated by state and federal agencies, with the Northeast Fisheries Science Centers (NEFSC) twice-annual bottom trawl survey the most expansive in spatial and temporal coverage. This survey has been a critical tool in monitoring the relative abundance of groundfish stocks from Cape Hatteras, NC to Nova Scotia since the 1960s. However, bottom trawl surveys like the NEFSCs have reduced efficiency over complex bottom habitats with high bathymetric relief or hard substrate, such as cobble fields or rocky ledges (McElroy et al. 2019; Grabowski et al. 2020). Most bottom trawl survey programs have low sampling effort within shallow or complex habitat areas due to risks to the equipment, and will instead focus on sampling in areas deeper than 18m and with soft and smooth bottom habitats (Johnston and Sosebee 2014).

The limited survey information from complex bottom habitats is concerning to fishing industry stakeholders, fisheries scientists, and fisheries managers alike. Habitat complexity likely interacts with catchability, thereby making it difficult to determine if differences in catch between habitats are truly reflective of relative abundance (Peterson and Black 1994; Grabowski et al. 2020). The challenge that this poses to cod stock assessments can vary by life history phase; several early life history phases of cod are associated with complex bottom habitat areas and therefore are not likely to be sampled well by conventional bottom trawl methods. Age-0 and age-1+ juvenile cod have been found in higher densities over hard substrate or high-bathymetric relief areas, likely as a refuge from predation (Gotceitas and Brown 1993; Gotceitas et al. 1995; Gregory and Anderson 1997; Cote et al. 2004; Lough 2010; Grabowski et al. 2018; Linner and Chen 2022). Though conventional wisdom holds that adult cod prefer colder and deeper offshore waters, recent evidence indicates that shallow inshore areas support a wide size range of cod (Dean et al. 2021). Industry stakeholders also have reported a relatively high density of large cod within inshore hard-bottom habitats of the western Gulf of Maine, possibly indicating a density-dependent reduction in large cod spatial distribution and altered availability to bottom trawl surveys (Grabowski et al. 2020; McElroy et al. 2021). Stakeholders observations of increased cod density over complex bottom habitats have created the perception that cod abundance across most of the species’ spatial range is much higher than what stock assessments have suggested, which has strained relationships between scientists, managers, and stakeholders.

A further complication to assessing cod population dynamics is the complex spatial structure of its subpopulations within the NEUS region. Since 1972, cod in this large marine ecosystem have been managed as two spatially distinct stock units: the Georges Bank and Gulf of Maine stocks (Serchuk and Wigley 1992). Cod within the far eastern section of Georges Bank are within the Canadian exclusive economic zone and are jointly managed by Canada and the US. This stock structure was based on the state of cod distribution and connectivity knowledge at the time, but more recent research indicated that the true biological structure of cod is more complex, leading to misinterpretations of the spatial variation and magnitude of cod productivity (Kerr et al. 2014; Zemeckis et al. 2014b). Recent synthesis by the Atlantic Cod Stock Structure Working Group provided evidence that there are five biological stocks of cod in US waters: Georges Bank, Southern New England, Eastern Gulf of Maine, and two sympatric Western Gulf of Maine stocks with distinct spawning times in winter and spring (McBride and Smedbol 2022). This synthesis was the impetus to adjust the spatial areas for assessment from the former two units (Gulf of Maine, Georges Bank) to four newly-defined units–Southern New England (SNE), Georges Bank (GBK), Eastern Gulf of Maine (EGOM), and Western Gulf of Maine (WGOM; spring and winter spawners combined) (Fig. 1). Each of these spatial areas has a unique composition of static spatial features (e.g., depth, bottom substrate, rugosity) and dynamic environmental characteristics (e.g., bottom water temperature) that are expected to influence the within-stratum distribution and productivity of the associated cod stock (Ames 2004; Zemeckis et al. 2014b; Guan et al. 2017a; Dean et al. 2019; Linner and Chen 2022). Tracking the varying spatial dynamics and productivity of cod stocks at spatially explicit and biologically relevant scales will be critical to informing accurate assessments and developing effective management strategies (Kerr et al. 2014; Zemeckis et al. 2014b; Dean et al. 2019; McBride and Smedbol 2022).

**Figure 1:**
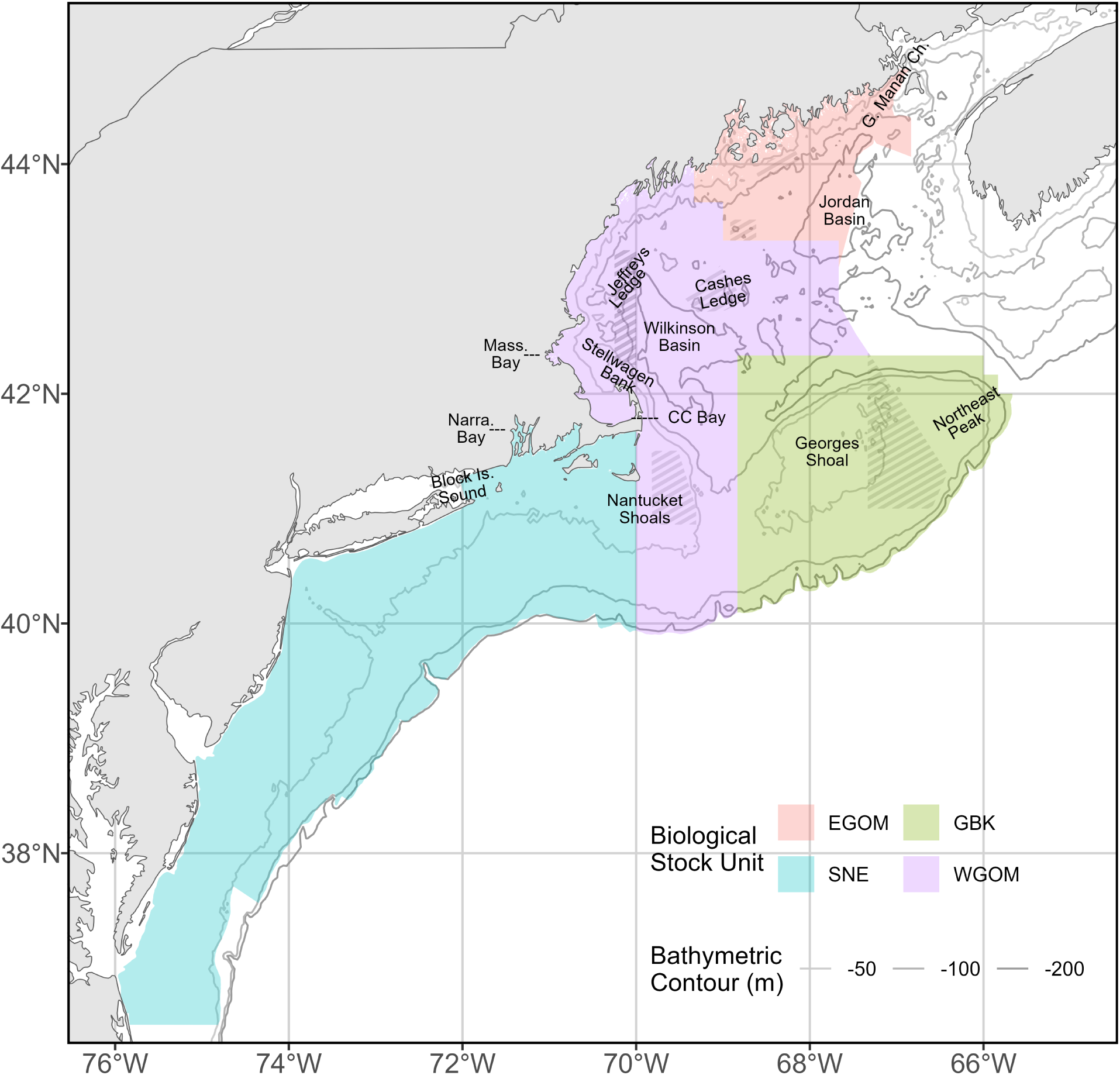
Spatial domain of cod VAST models with biological stock spatial strata (EGOM: Eastern Gulf of Maine, GBK: Georges Bank, SNE: Southern New England, WGOM: Western Gulf of Maine). Areas with year-round groundfishing closures are filled with crosshatches. Lines indicate the 50, 100, and 200 m isobaths, with darker lines representing deeper isobaths. Ecologically important regions are annotated on the map.

The objective of this study was to build size-specific and seasonal maps of spatial density for Atlantic cod for each of the stock areas using all relevant state and federal groundfish survey data. Surveys available for inclusion in modeling cover inshore, offshore, smooth, and complex bottom habitats, and utilize both bottom trawl and bottom longline methods. Vector Autoregressive Spatio-Temporal (VAST) models were used to combine all survey data into size-specific models of cod spatiotemporal density. VAST models estimate the spatiotemporal density of a target organism conditioned on density covariates and controlling for catchability covariates, and so are useful for estimating the spatiotemporal density of organisms in areas with limited observation data. These estimates of spatial-temporal density can then be used in the calculation of indices of relative abundance, centers of gravity, range edges, and habitat associations. VAST models were fitted for three size classes of cod to examine whether cod size influences distribution, abundance, and habitat use patterns. The output of these models can inform our understanding of cod spatial dynamics, habitat associations, and demographics, benefiting management efforts.

## 2 Methods

### 2.1 Survey Data

Eleven surveys of groundfish abundance were available for use (Table 1). The combined spatial footprint of all surveys covered the waters of the continental shelf along the US coast from Lubec, Maine to Cape Hatteras, North Carolina. Temporal coverage spanned from 1959 to 2022. These surveys utilized different vessels and methods, have variable spatial extents, were completed either annually or seasonally, and had variable temporal coverage (as in, the number of years in which the survey occurred). All surveys reported the number, individual lengths, and aggregate weight (kg) of cod caught per tow, and some surveys processed all or a portion of the catch to provide further biological detail (individual weight, sex, age, etc.). Survey data were cleaned (see next section), cut to chosen model spatial and temporal domains, and combined into a single dataset. Leave-one-out sensitivity tests were used to select which survey data would be included in the modeling process (see supplemental material).

**Table 1:** Description of the 11 surveys available to model groundfish spatiotemporal distribution. Included for each survey are details for gear type, cod biological stock areas covered by the survey (EGOM: Eastern Gulf of Maine, GBK: Georges Bank, SNE: Southern New England, WGOM: Western Gulf of Maine), years in which the survey was conducted, seasons in which the survey is conducted, and method of calculating area swept.

| Survey | Abbreviation | Gear | Stocks | Years | Seasons | Area Swept |
| --- | --- | --- | --- | --- | --- | --- |
| Northeast Fisheries Science Center Bottom Trawl | NEFSC BTS | Bottom trawl | EGOM, GBK, SNE, WGOM | 1963-2023 | Fall, Spring | Average reported |
| Canadian Department of Fisheries and Oceans Bottom Trawl | DFO Trawl | Bottom trawl | GBK | 1986-2023 | Spring, Summer | Average reported |
| Maine-New Hampshire Inshore Bottom Trawl | ME-NH Trawl | Bottom trawl | EGOM, WGOM | 2000-2023 | Fall, Spring | Average reported |
| Massachusetts Division of Marine Fisheries Inshore Bottom Trawl | MADMF Inshore Trawl | Bottom trawl | WGOM, SNE | 1978-2023 | Fall | Average calculated |
| Massachusetts Division of Marine Fisheries Industry-Based Trawl | MADMF Industry | Bottom trawl | WGOM, SNE | 2003-2007; 2016-2019 | Fall, Spring | Exact reported per haul |
| Atlantic States Marine Fisheries Commission Northern Shrimp Trawl | ASMFC Shrimp Trawl | Bottom trawl | WGOM, SNE | 1983-2023 | Summer | Average calculated |
| Rhode Island Department of Environmental Management Coastal Trawl | RIDEM Trawl | Bottom trawl | SNE | 1979-2023; 1990-2023 | Fall, Spring; Monthly | Average calculated |
| University of Rhode Island Graduate School of Oceanography Fish Trawl | URI GSO Trawl | Bottom trawl | SNE | 1959-2023 | Weekly | Average calculated |
| UMass Dartmouth School for Marine Science & Technology Video Trawl | SMAST Trawl | Bottom trawl with camera | WGOM, SNE | 2016-2022 | Winter, Fall, Spring | Exact reported per haul |
| Cooperative Gulf of Maine Bottom Longline | NEFSC BLLS | Bottom longline | WGOM, SNE | 2014-2023 | Fall, Spring | Undefined, assumed |
| Eastern Gulf of Maine Sentinel | Sentinel | Jigging | EGOM | 2012-2022 | Fall | Undefined |

#### 2.1.1 Data cleaning and response variable

To be considered in the model, survey data needed to provide valid information for spatial location, sampling date, and number of cod caught, as well as valid size-identifying biological information. Cod caught in the surveys were separated into three distinct size classes, as it was expected that habitat utilization and spatial density would vary among size or age groups. Small cod were defined as shorter than 39.1 cm total length or less than 0.58 kg. Medium cod were between 39.1 and 70.2 cm, or between 0.58 and 3.44 kg. Large cod were longer than 70.2 cm or heavier than 3.44 kg. This size structure roughly matches ages 0-2 (pre-spawning), ages 2-5 (variable spawning proportions at age), and ages 5+ (spawning) cod (Zemeckis et al. 2014b; Dean et al. 2019; Dean and Perretti 2022). This size-based classification structure and its general relationship to maturity is assumed to be consistent across all years and all spatial areas of the model domains, though time to maturity may be slightly different between time periods and spatial locations (Dean and Perretti 2022). Fish that were unable to be assigned to a size class were not used in the models. Because most surveys only collect individual weight information for a subsample of total catch, biomass per size class could not be used as the modeled response variable. Instead, total abundance (count) of each of the size classes was used as the response variable for each sampling event.

### 2.2 VAST

VAST models were used to estimate cod spatial density over time by size class and create joint indices of relative abundance using the cleaned groundfish survey data. VAST is a framework for implementing spatial delta-generalized linear mixed models (delta-GLMM) and can be structured to provide estimates for multiple categories of interest and spatial strata (Thorson and Barnett 2017; Thorson 2019). As recommended by model developers for abundance-data models, we specified a Poisson-link delta model with lognormal-Poisson distribution. This model family is designed to accept positive continuous data with zeros and use a delta (or hurdle) approach to derive spatiotemporal density as a combination of two linear predictors (Thorson 2018). The first linear predictor estimates encounter probability, and the second linear predictor estimates catch rates given a positive encounter probability. Using notation from Thorson (2019), the first linear predictor can be represented as

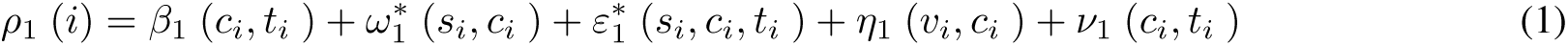

where *ρ*_1_ (*i*) is the encounter probability predictor for observation *i* for category *c_i_* at location *s_i_* and time *t_i_*. *β*_1_ (*c_i_, t_i_*) represents temporal variation for each category and time, *ω_i_* (*s_i_, c_i_*) represents spatial variation for each location and category, *ε*_1_ (*s_i_, c_i_, t_i_*) represents spatiotemporal variation for each location, category, and time, *η*_1_ (*v_i_, c_i_*) represents vessel effects for each vessel and category, and *ν*_1_ (*c_i_, t_i_*) represents the effect of density covariates for each category and time. The second linear predictor is structured the same way but calculates the catch rate predictor. Both linear predictors incorporate fixed and random effects, and spatial and spatiotemporal variation are approximated as Gaussian Markov random fields (Thorson et al. 2015b; Thorson and Barnett 2017; Thorson 2019).. Implementation of VAST models requires several further structural and data inclusion decisions, as outlined in Thorson (2019) and below in sections 3.2.1 through 3.2.6.

Recent works have highlighted the sensitivity of indices of abundance derived from spatio-temporal delta-models models like VAST to a number of structuring decisions (Thorson et al. 2021; Cacciapaglia et al. 2024). When absolute abundance is the desired output of a delta-model, additional model structuring decisions and model selection processes are recommended, as is a comparison to design-based indices of abundance. However, the goal of this study was not to derive absolute abundance. Therefore, no quantitative comparisons between model-based and design-based indices of abundance were made, and indices of relative abundance presented in this study should not be used as representative of absolute regional abundance.

#### 2.2.1 Spatial domain, smoothing, resolution, and strata

Though survey data coverage extends to Cape Hatteras, North Carolina, observations of cod are rare further south than Delaware Bay. Therefore, the spatial extent of the model domain was US waters on the continental shelf from the northern edge of the Gulf of Maine through the mouth of the Chesapeake Bay, with extension into Canadian waters for the jointly-managed eastern Georges Bank area (Fig. 1). This area was represented as a 2D mesh built on a stochastic partial differential equation (SPDE) approximation to a Gaussian Markov random field with a Matérn correlation function. Geometric anisotropy (directional correlation) is expected in most marine ecosystems (Thorson et al. 2015a) and was therefore included as a fixed effect, though support for its inclusion was also assessed in the model selection process (see Section 3.3). Spatial variables were defined at a pre-determined number of knots. Knots were placed via k-means clustering of the data to minimize the average distance between knots and sampling locations. Sampling locations are expected to have spatial variables equal to the nearest knot, so in effect, the number of knots defines the spatial resolution of spatial density estimates. In this model, the number of knots was set to 200 and the mean distance between nearest-neighbor knot locations was 30.1 km. Assessment of directional correlation found that the distance with approximately 10% correlation was 123.2-157.7 km for the first linear predictor and 42.7-66.0 km for the second linear predictor across all size classes, indicating that this number of knots and the resulting distance between knots provided sufficient spatial resolution.

It is of interest to calculate spatial dynamics and indices of relative abundance for each of the four biological stock areas proposed by the Atlantic Cod Stock Structure Working Group, as there is evidence that the complex spatial structure of these biological stocks affects both our understanding of cod spatial dynamics and management efforts (Zemeckis et al. 2014b; Guan et al. 2017a; McBride and Smedbol 2022; Linner and Chen 2022). Therefore, a custom extrapolation grid was built as a spatial domain for derived quantities, which had 2000 grid cells (each cell approximately 20.1 km by 20.1 km) spread across four spatially defined stock areas. Bilinear interpolation was then used to calculate spatial density within each of these cells. The spatial strata represented the Eastern Gulf of Maine, Georges Bank, Southern New England, and combined winter-and spring-spawning Western Gulf of Maine stocks. The latter two stocks overlapped substantially in space, supporting their treatment as a single spatial stratum. The models also reported results at a basin-wide scale, in which all strata were combined.

It is of interest to calculate spatial dynamics and indices of relative abundance for each of the four biological stock areas proposed by the Atlantic Cod Stock Structure Working Group, as there is evidence that the complex spatial structure of these biological stocks affects both our understanding of cod spatial dynamics and management efforts (Zemeckis et al. 2014b; Guan et al. 2017a; McBride and Smedbol 2022; Linner and Chen 2022). Therefore, a custom extrapolation grid was built as a spatial domain for derived quantities, which had 2000 grid cells (each cell approximately 20.1 km by 20.1 km) spread across four spatially defined stock areas. Bilinear interpolation was then used to calculate spatial density within each of these cells. The spatial strata represented the Eastern Gulf of Maine, Georges Bank, Southern New England, and combined winter-and spring-spawning Western Gulf of Maine stocks. The latter two stocks overlapped substantially in space, supporting their treatment as a single spatial stratum. The models also reported results at a basin-wide scale, in which all strata were combined.

#### 2.2.2 Temporal domain and resolution

The temporal domain of the models began in 1982 when the adoption of the Interim Groundfish Plan shifted management strategies from regional quotas to minimum size and gear regulations. The last full time step of data available at the time of modeling was fall 2021. Survey data from outside this temporal domain were not used in the model.

Many surveys considered by this modeling effort were conducted twice annually, in the spring (March-May) and fall (September-October). This was thought to be a useful sampling design to track seasonal migrations of marine fishes. In the highly-studied western Gulf of Maine region, spring-spawning cod typically migrate inshore to spawn April to July, move offshore to feeding areas in the summer and fall, and may move to deep offshore basins to overwinter (Zemeckis et al. 2017). Winter-spawning cod in the same area will move inshore to spawn November to December (Zemeckis et al. 2014b). Across all biological stock areas, cod exhibit spawning site fidelity and make seasonal migrations to these areas (Ames 1997, 2004; Kovach et al. 2010; Zemeckis et al. 2014b, 2014a; DeCelles et al. 2017; Clucas et al. 2019; McBride and Smedbol 2022). Because it is supported both by data availability and the behavior of cod, time steps in the model were structured to represent the spring and fall seasons of each year in the time series. Therefore, though there were 40 years of data, there were 80 time steps. The spring season was March through August of any year *x*, and Fall was September through December of year *x* and January and February of year *x*+1. This approach ensured that the fall season time steps were temporally continuous. Therefore, day 1 of a modeled year is March 1^st^.

#### 2.2.3 Effort estimates

VAST requires an effort estimate for each observation. For surveys using bottom trawl methods, area swept is a commonly reported effort measure. Area swept was reported, assumed, or calculated for the surveys included in this modeling effort as information was available (Table 1). Some bottom trawl surveys reported area swept for each tow, and this was therefore used as an effort measure. Several surveys did not report the area swept for each tow but instead reported an average area swept based on gear mensuration and vessel travel distance. Typically, these surveys also validated that effort was within tolerance limits for acceptable tow duration and vessel speed to maintain similarity between tows. For these surveys, this provided average area swept was included as the estimated effort for each observation. Finally, a few surveys reported only optimal gear mensuration and intended distance towed. For these surveys, the estimated average effort per tow was calculated as the intended distance covered by the tow multiplied by the optimal wing spread.

It is recommended that the area swept be set to 1 for sampling gears with an unknown effective area swept (Thorson 2019). Initially, the area swept for both the Eastern Gulf of Maine Sentinel survey and the bottom longline survey was set to 1. However, this created an issue of scaling and mixed units. The description of the bottom longline survey motivation and methods in McElroy et al. (2019) states that it was developed to match the sampling effort of the NEFSC bottom trawl survey as closely as possible. The bottom longline survey uses a 1 nautical mile groundline soaked for 2 hours across slack tide in an attempt to approximate the same sampling area of the NEFSC bottom trawl survey. The two surveys caught comparable numbers of cod per unit effort across all size classes and seasons (Fig. S1). For these reasons, the average area swept of the NEFSC bottom trawl survey was used as the input for the area sampled for each set of the bottom longline survey. It is likely that the true area sampled of the bottom longline survey varies with current velocity and other oceanographic conditions, but a longer time series and further calibration studies are needed to make a true quantitative determination. There is no estimation or accepted assumption of area sampled for jigging surveys like the Sentinel survey, and therefore it had to be excluded. This removed a total of 332 samples between 2012 and 2021.

#### 2.2.4 Spatial, temporal, and spatiotemporal effects

Spatial, temporal, and spatio-temporal effects can be included in both linear predictors. A model selection process was used to justify the use of spatial and spatiotemporal random effects in the first and second linear predictors. The intercept for each linear predictor was defined as a fixed effect for each time step this ensured independent estimates of abundance for each time step, which is most appropriate for creating abundance indices (Thorson 2019). A temporal correlation component was estimated for the spatio-temporal variation in both linear predictors. This is recommended for indices generated by multiple data sources that do not necessarily sample the same locations in every time step (Thorson 2019). Without this estimation, unrealistic hot spots may develop or be carried through the time series when this is inappropriate. Because the models include data from surveys with varied sampling intensity, locations, and temporal coverage, this is an important structuring decision.

The model used to estimate the temporal correlation component varied with size class. VAST models for small-and medium-sized cod included sufficient observation data to successfully fit an AR1 process. Attempts to use an AR1 process for the large size class failed to fit. A random walk process was used instead. For all size classes, the temporal correlation component was calculated for the first linear predictor. These results were used as the temporal correlation components of the second linear predictor, rather than calculating a new temporal correlation component for the second linear predictor.

#### 2.2.5 Vessel effects

The random variation in catchability among levels of a grouping variable is referred to as vessel effects in the VAST model structure. VAST models covariation in vessel effects with a factor model, where variation in catchability between groups is a random effect. Each survey used in this modeling effort has its own set of sampling protocols and vessels, and these differences are likely to introduce variability in catchability. Therefore, vessel effects were included in the models using survey as the grouping variable.

#### 2.2.6 Density Covariates

VAST allows for the effects of both density and catchability covariates to be included in modeling efforts. Catchability covariates are processes that affect the ability to observe the target organism without necessarily affecting the distribution of the organism. Density covariates are processes that directly affect the distribution of the target organism, regardless of the ability to observe it. Both covariates affect the catch rate of the target organism, but only density covariates are used to predict target organism density within the spatial domain. Therefore, VAST controls for catchability covariates and conditions on density covariates. VAST is unable to distinguish whether potential covariates should be treated as catchability or density covariates; this must be decided with theoretical insight from an analyst. As mentioned previously, differences in sampling design were included as vessel effects, but explicit catchability covariates were not used.

##### Bathymetry

Several environmental variables were tested as potential density covariates. The final model only includes density covariates with significant impact, as determined by a model selection process outlined in Section 3.3. Depth strongly influences the distribution and habitat use of Atlantic cod (Lough 2010; Guan et al. 2017b; Li et al. 2018; Linner and Chen 2022). Very few cod are found in waters deeper than 400 m, and the highest densities of cod are found between 10 and 150 m (Lough 2010). Depth at all survey locations was extracted from rasterized GEBCO 15 arc-second bathymetry (Group 2023) and included as a potential density covariate.

##### Sediments

There is evidence that cod habitat preferences include large-grain sediments like gravel, cobble, and boulders, making sediment type an important environmental covariate to consider when mapping spatial density (Gotceitas and Brown 1993; Gotceitas et al. 1995; Methratta and Link 2006; Lough 2010; Grabowski et al. 2018; Linner and Chen 2022). The spatial distribution of sediment types through the VAST models spatial domain was modeled by Brad Harris and Felipe Restrepo at Alaska Pacific University (Harris and Restrepo, pers. comm.). This model is an expansion of the New England Fishery Management Council Swept Area Seabed Impact (SASI) model, which used sediment observations from many sources to model and classify bottom habitats by sediment particle size (Bachman et al. 2011, 2019). The sediment classes (mud, sand, gravel, cobble, and rock) were based on Wentworth (1922) (Table S1). The sediment distribution model used an ordinary kriging approach to interpolate the probability of finding any of the five sediment classes within the cells of a 1 km by 1 km resolution grid with the same spatial extent as the VAST model.

##### Rugosity

There is further evidence that cod prefer habitats with high bathymetric relief, like boulders and steep ledges (Gregory and Anderson 1997; Cote et al. 2004). Bathymetric relief was characterized by rugosity, which is a unitless measure of bottom vertical change over horizontal distance. Using methods outlined in Friedman et al. (2012), rugosity was calculated from the 15 arc-second rasterized bathymetry data over the VAST models spatial domain. Rugosity at each survey location was extracted from the resulting rugosity raster.

##### Bottom temperature

The previous density covariates are spatially dynamic but temporally stationary. Cod distribution is also often temporally dynamic, as cod have seasonal migrations that likely reflect shifting water temperatures (Lough 2010; Zemeckis et al. 2017; Li et al. 2018). Bottom water temperatures have been linked to cod productivity, and therefore likely affect cod distribution (Drinkwater 2005; Methratta and Link 2006, 2007; Guan et al. 2017b). Models of bottom water temperatures within the NEUS continental shelf were provided by Du Pontavice et al. (2023). The bottom temperature within approximately 5-minute by 5-minute grid cells was calculated at a daily timestep for 1982 to 2020. Bottom temperature data were not available for 2021, the final year in the VAST models temporal domain. Instead, bottom temperature data from 2020 were used to fill this gap, as it was assumed that bottom temperature trends would remain similar between sequential years. Further, the bottom temperature product did not extend to the inshore areas within the modeled spatial domain. To resolve this limitation, bottom temperature was extrapolated to the shoreline using an ordinary kriging approach.

##### Climate indices

Several climate indices have a relationship to cod distribution and abundance, as they are associated with long-term warming trends and likely reflect regional habitat suitability for cod (Pershing et al. 2015). For this model, North Atlantic Oscillation (NAO) and Atlantic Multidecadal Oscillation (AMO) index data were used as spatially static basin-wide climate indices. NAO is a measure of differences in atmospheric pressure at high and low latitudes of the North Atlantic and is expected to affect the intensity and location of wind patterns, heat transport, and moisture transport. AMO measures average anomalies of sea surface temperature in the North Atlantic basin. Data for both climate indices were publicly available in NOAAs data repositories. Values for both climate indices were extracted for every survey observation at the best available temporal resolution, which was a daily timestep for NAO and a monthly timestep for AMO.

### 2.3 Model selection

The model selection process was conducted separately for each size class and consisted of two steps. The first step compared models with and without anisotropy and/ or spatial and spatiotemporal random effects in the linear predictors as in Ng et al. (2021) and Gaichas et al. (2023). These models did not include any density covariates. AIC was used to compare models, and restricted maximum likelihood (REML) was used in model construction to make comparison via AIC possible (Zuur et al. 2009). For all three size classes, the best model included anisotropy and spatial and spatiotemporal effects in both linear predictors (Table S2).

Before inclusion in model selection, potential density covariates were tested for collinearity. If collinear pairs were found, only the most ecologically relevant density covariate was selected for inclusion (see supplemental material). The second step in model selection was to select the most informative combination of potential density covariates. For each size class, a series of drop-one covariate models were run to assess covariate influence via AIC, as in Hansell et al. (2022). If AIC values were within 2 units of each other, the most parsimonious model was selected (Burnham and Anderson 2004) (Table S3).

### 2.4 Final model, diagnostics, and derived quantities

After the model selection process, the best selected models for all three size classes were run twice; once with Gaussian Markov Random Fields (GMRF) for enhanced fine-scale spatiotemporal interpolation turned on, and once with them turned off. The resulting centers of gravity from model fits were compared. Major differences between centers of gravity from models with and without GMRF turned on would indicate an important density covariate is not being explicitly modeled (Perretti and Thorson 2019; Hansell et al. 2022).

Final models were run with both GMRF and bias correction turned on. Models with bias correction use the epsilon method to ensure that the mean and variation of generated indices of relative abundance are not biased due to their transformation by a nonlinear function in the modeling process (Thorson and Kristensen 2016). Mapping residuals within the spatial domain did not highlight any spatial area as having a consistently poor fit (Fig. S2-S4)

After final model selection and assessment of diagnostics, seasonal maps of cod spatial density were generated for each size class. These maps are useful on their own to visualize changes in cod spatial density through space and time but can also be used to derive other measures of population and spatial dynamics. Resulting maps were used to derive spring and fall indices of relative abundance for all three size classes of cod, both for individual biological stock areas and for the entire NEUS region. Seasonal centers of gravity (COG) were also derived for each biological stock area and for the overall NEUS region. COG measures the density-weighted spatial location of the center of a selected population. Range shifts were quantified by the derivation of seasonal northeastern and southwestern range edges, represented by the 0.05 and 0.95 quantiles of cod distribution along the northing and easting axes, though these could only be calculated at the spatial scale of the entire modeled domain. It should be noted that the northeastern range edge is not a true quantification of the northern or eastern limits of the northwest Atlantic cod population, as the modeled spatial domain excludes significant cod habitats in Canadian waters. The distance between seasonal range edges along a directional axis (e.g., north-south, east-west) was calculated and used alongside range shift metrics as a measure of habitat compression or expansion. Together, these results could indicate whether changes in cod abundance are more likely due to spatial changes in productivity within a consistent range or range shifts. Finally, habitat associations were quantified via conditional response plots. Though these conditional response plots could only be derived for the entire modeled domain and without seasonal distinction, they helped identify which density covariates had the most influence on cod encounter and catch rates.

## 3 Results

For all size classes, the effect of anisotropy was stronger in the first linear predictor than the second, meaning that the encounter rate was similar for longer distances along a directional axis than the positive catch rate. The axis of anisotropy for all size classes generally ran southwest-to-northeast, indicating a greater degree of similarity along the NEUS coastline than along an inshore-offshore gradient (Fig. S5-S8).

### 3.1 Small cod

The series of drop-one models to test for density covariate significance indicated that monthly AMO was not a significant predictor of small cod density, and so it was excluded from all small cod model runs. Small cod densities were highest in nearshore waters along the coast from Massachusetts Bay to Narragansett Bay, particularly in the spring time series (Fig. S6a). The spring index of relative abundance had high interannual variability and a sudden increase in 2003 but consistently indicated that the WGOM and SNE stocks contributed the most to NEUS regional abundance (Fig. 2a). Spring COG for all small cod within the NEUS region was typically located around Cape Cod, on the border of the WGOM and GBK stocks, and was highly variable throughout the time series with no clear directional trend (Fig. 3a). Stock-specific COGs indicated that changes in the distribution of cod within the WGOM stock had the most influence on the northing component of regional COG, but changes in relative abundance at the westernmost-known spawning and settlement locations (west of Marthas Vineyard, SNE biological stock area) likely influenced the easting component of regional COG (Fig. 4a, S6a). Similarly, spring range edges were highly variable (Fig. 5a, g). The southwestern range edge, represented by the 0.05 quantile of density-weighted distribution, has remained around the same north-south location since the 1990s, but oscillated east-west up to 85 km per year as the highest density patches shifted between Block Island Sound and Cape Cod Bay. The northeastern range edge, represented by the 0.95 quantile of density-weighted distribution, had rapid annual north-south and east-west shifts as relative productivity between the EGOM, WGOM, and SNE stocks changed. The distance between spring southern and northern range edges had high interannual variability from 2002 onwards, with evidence of north-south range compression beginning in 2016 as the northern range edge shifted south. The distance between the eastern and western range edges compressed by approximately 62 km from 1982 to 2002 as all stocks but WGOM saw decreases in productivity. Since 2002, increased productivity near Cape Cod has led to an approximately 17 km increase in east-west range.

**Figure 2:**
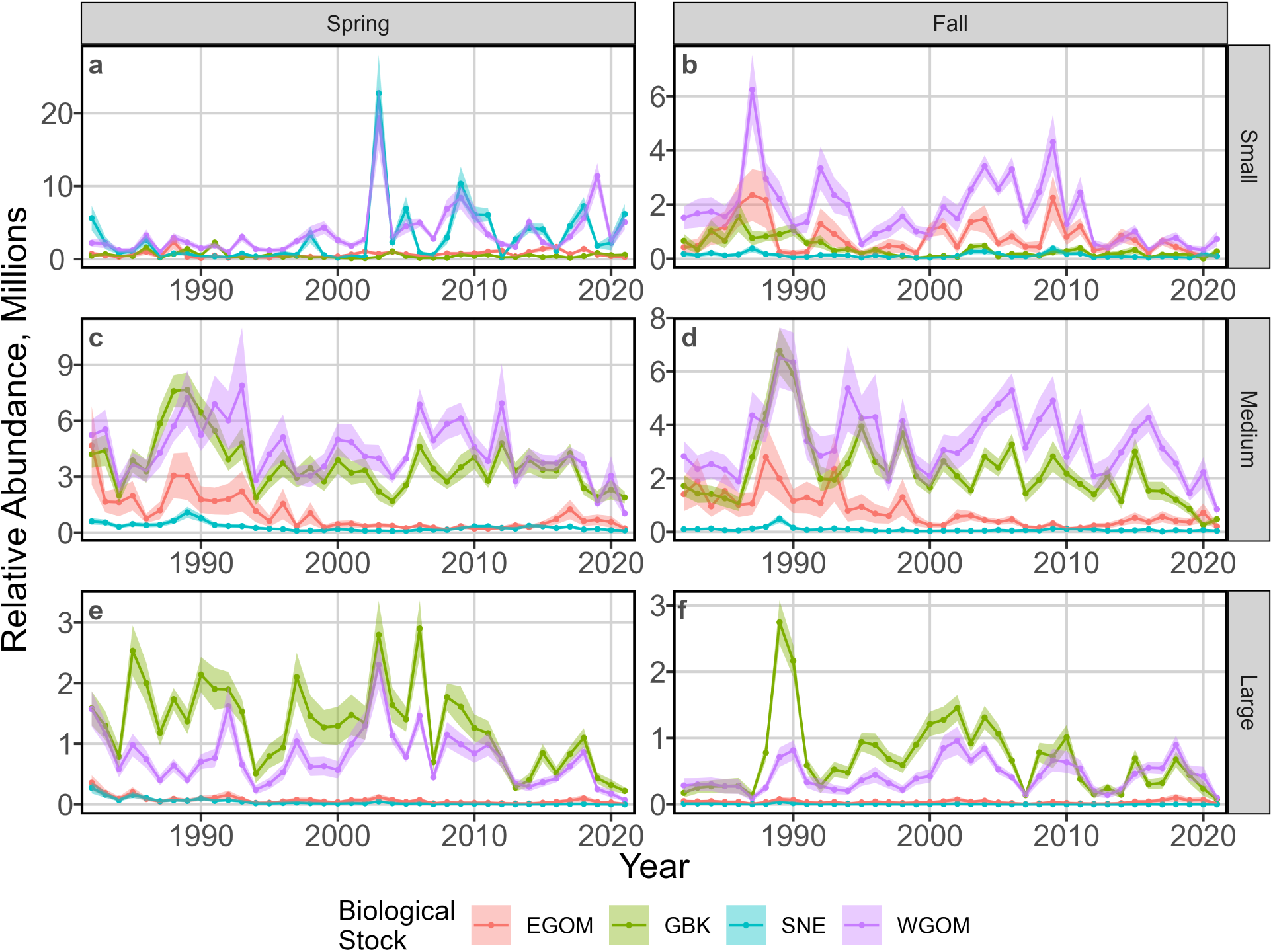
Indices of relative abundance derived from model outputs. Results are reported in millions and separated by season (columns) and size class (rows).

**Figure 3:**
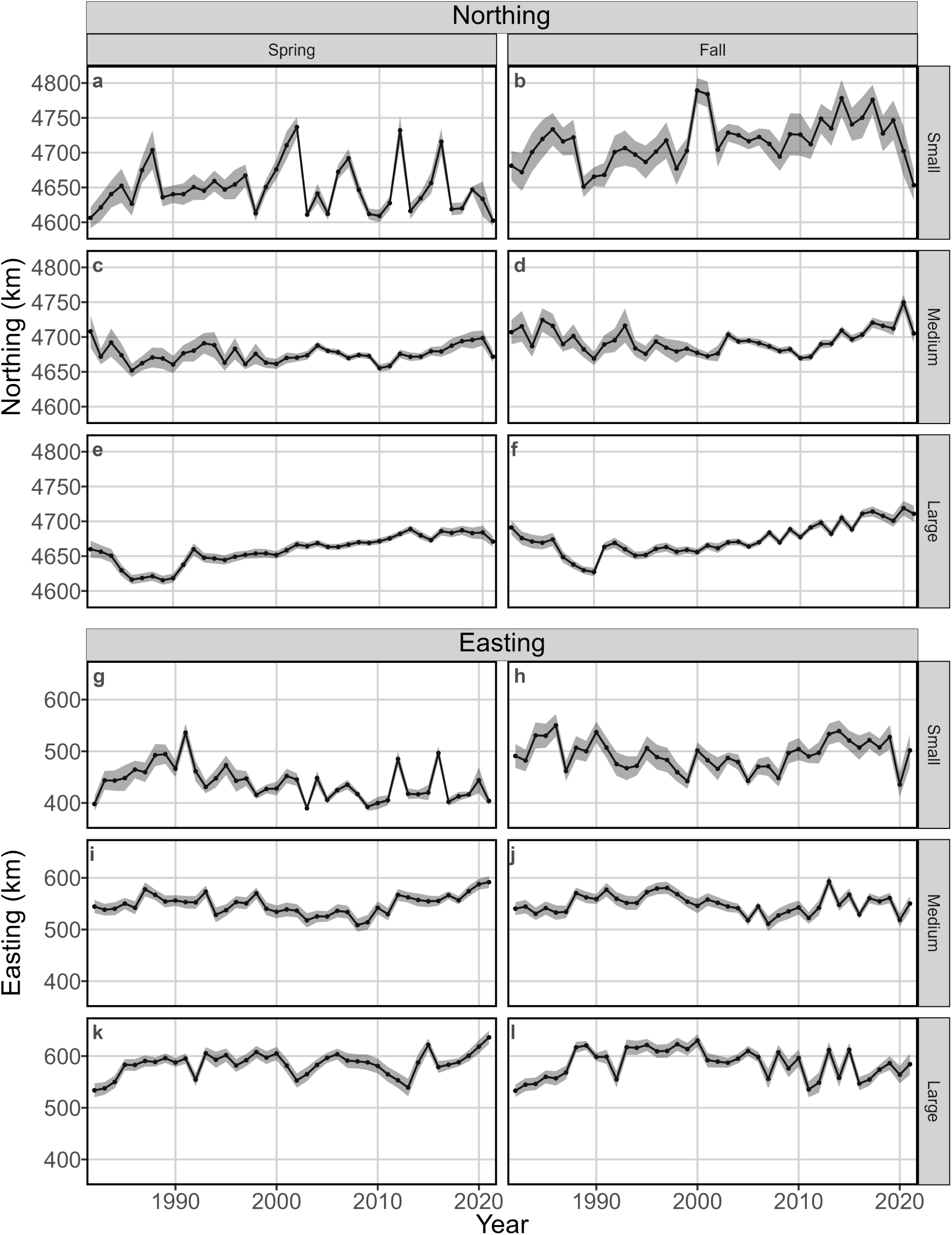
Population centers of gravity over the time series. Population centers of gravity represent the center of cod size class-and season-specific density surfaces without considering the boundaries of biological stock areas. Results were calculated using a WGS84 projection and UTM Zone 19N coordinate reference system. Eastings are therefore reported as kilometers from the western edge of the reference frame, and northings as kilometers from the southern edge of the reference frame. Results are separated by season (columns) and size class (rows). Northings are reported in frames a-f and eastings are reported in frames g-l.

**Figure 4:**
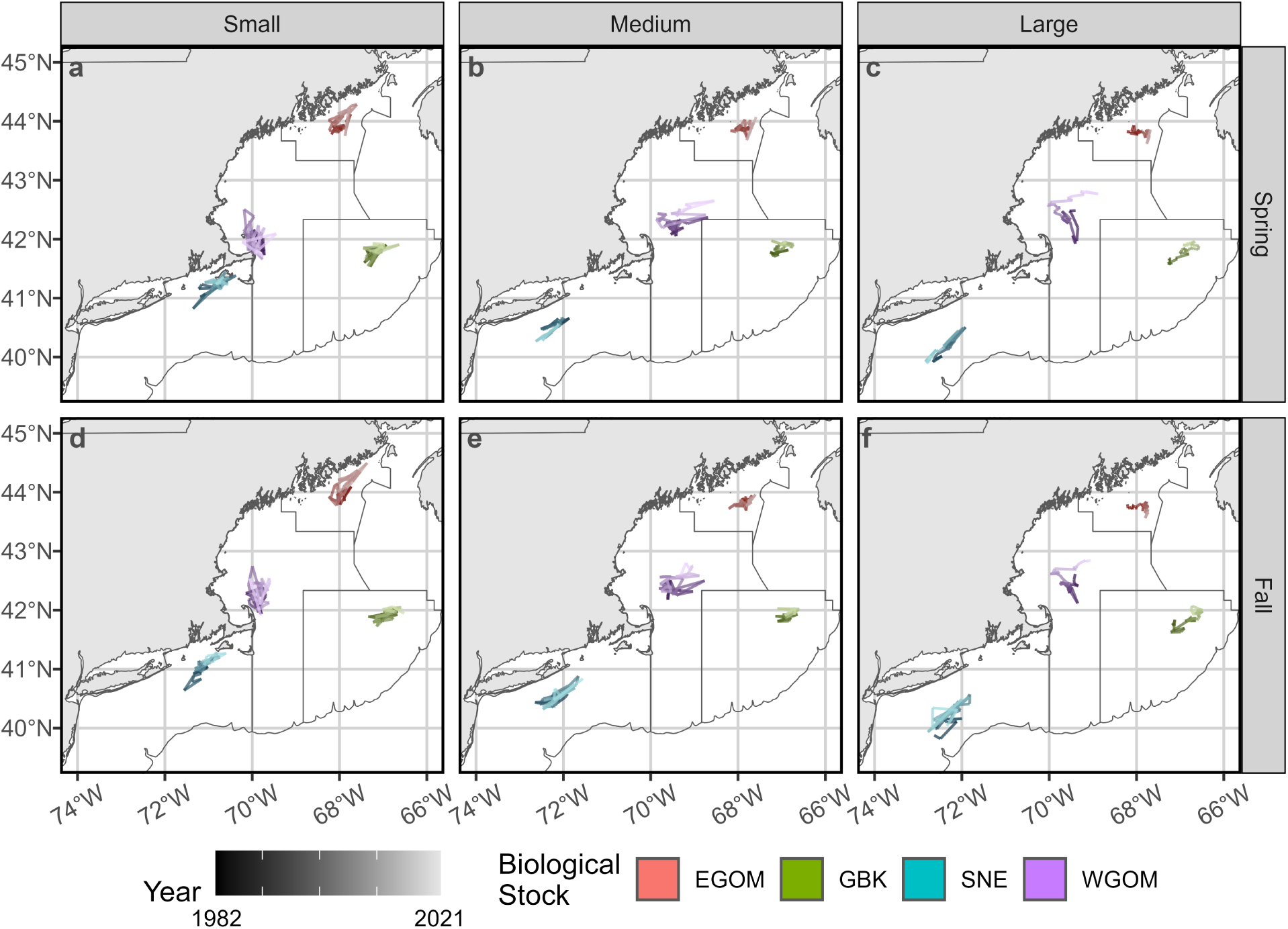
Spatial location of stock centers of gravity within the modeled domain. Maps were created using a WGS84 projection and UTM Zone 19N coordinate reference system. Stock area boundaries are represented by black lines; see figure 1 for more detail on each stock area. Results are separated by size class (columns) and season (rows). The southern portion of the Southern New England biological stock area is not represented so that more detail can be seen in center of gravity shifts. Darker, more saturated colors represent stock centers of gravity earlier in the time series and lighter, less saturated colors represent stock centers of gravity later in the time series. Red gradients represent the Eastern Gulf of Maine (EGOM), greens represent Georges Bank (GBK), blues represent Southern New England (SNE), and purples represent the Western Gulf of Maine (WGOM).

**Figure 5:**
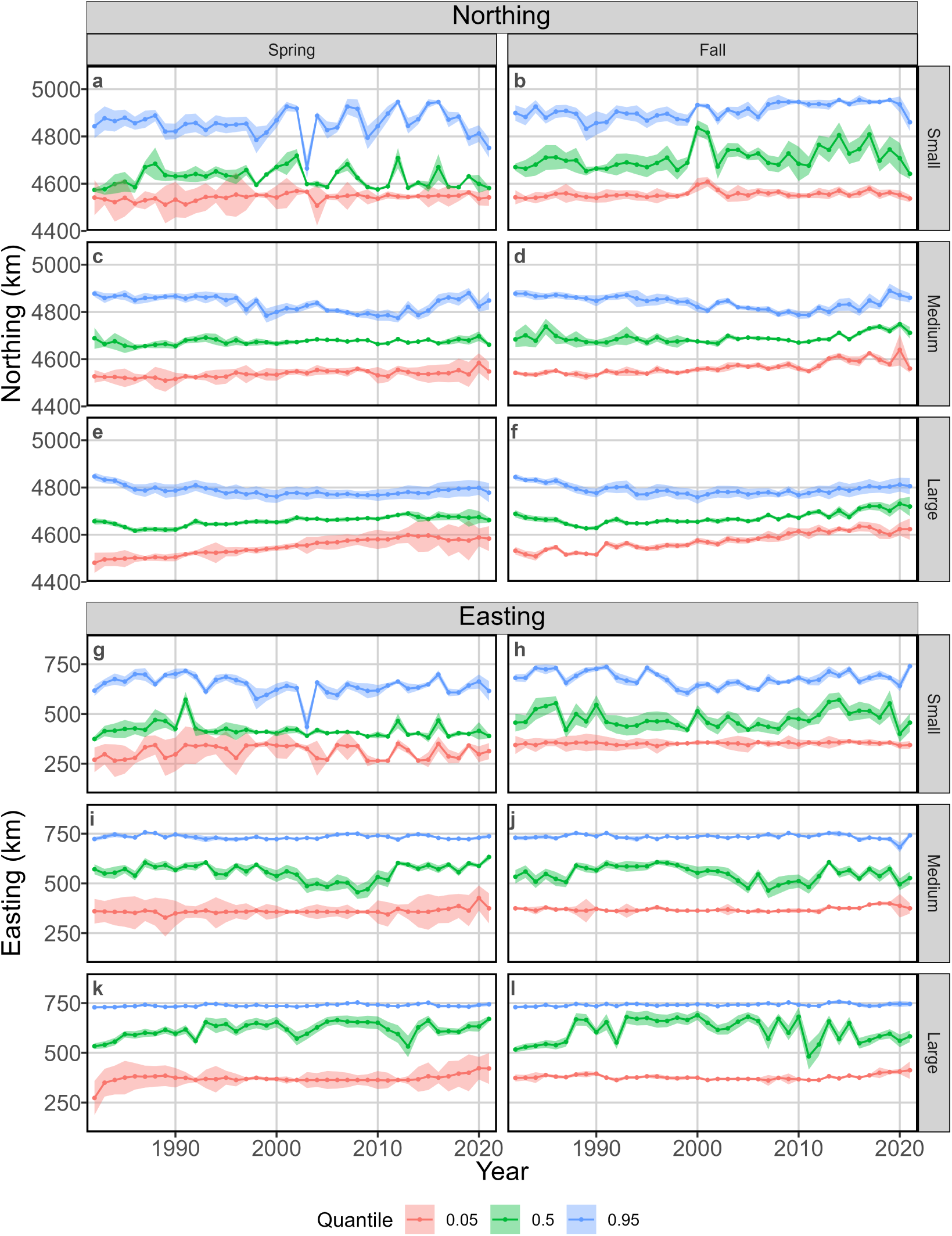
Population range edges over the time series. Results were calculated using a WGS84 projection and UTM Zone 19N coordinate reference system. Eastings are therefore reported as kilometers from the western edge of the reference frame, and northings as kilometers from the southern edge of the reference frame. Results are separated by season (columns) and size class (rows). Northings are reported in frames a-f and eastings are reported in frames g-l.

In the fall time series, small cod were present at highest densities in nearshore waters from Massachusetts Bay to the southern extent of Nantucket Shoals (Fig. S6b). Temporally inconsistent pockets of high density sometimes occurred at Cashes Ledge and in the waters between Grand Manan Island and Cutler, Maine. The fall index of relative abundance had two cycles of increase and decrease over the modeled periodabundance increased from 1982 to 1987, decreased until 1996, increased again until 2009, then rapidly decreased to the present day (Fig. 2b). WGOM and EGOM stocks followed these trends and consistently contributed the most to regional abundance, though the decline in the WGOM stock in the final 10 years of the time series was much faster than the decline in the EGOM stock. GBK and EGOM stocks contained a similar proportion of regional abundance at the beginning of the time series, but abundance in GBK declined in the 1990s and has remained low since then. Fall regional COG was typically located in the western Gulf of Maine, with a clear northward shift over time (Fig. 3b, h). When calculated for the entire modeled spatial domain, regional COG shifted on average 1.2 km/ year north. The east-west movement of the fall regional COG was less clear, with a period of westward (inshore) movement between 1982-2008, then a rapid return eastward (offshore) from 2009-2019. This likely reflects changes in relative productivity between stock areasregional COG was farther west when the relative proportion of regional abundance within the EGOM biological stock area was low, and farther east when the EGOM stock area contained proportionally more of regional abundance (Fig. 2b, Fig. 3b, h). Most stock COGs shifted north and east over the time series (Fig. 4d). Fall southwestern range edges remained in approximately the same place, while northeastern range edges have shifted north and east since the late 1990s as the EGOM stock has contained proportionally more of the regional abundance (Fig. 5b, h). The distance between the north and south range edges decreased by approximately 47 km from 1982 through 1999, and the distance between the east and west range edges decreased by approximately 83 km over the same period. Since 1999, the distance between north-south range edges increased by 13 km and the distance between the east-west range edges increased by 142 km. These distinct periods of range edge compression and expansion match with periods of GBK and EGOM stock decline (1982-1999) and slight recovery in the EGOM stock (2000-2021).

### 3.2 Medium cod

For the medium size class, all density covariates were retained as significant predictors. Models without AMO, bottom temperature, or predicted likelihood of cobble density covariates did not converge, highlighting their importance to modeling the spatial density of medium cod. Model outputs for the spring time series indicated medium-sized cod were consistently present at relatively high densities in all but the deepest sections of the Gulf of Maine and Georges Bank (Fig. S7a). Notable patches of high density include Nantucket Shoals, Stellwagen Bank and Jeffreys Ledge, Cashes Ledge, and the Northeast Peak region of Georges Bank. The index of regional relative abundance remained at relatively consistent levels between 1994-2018 but declined to a time-series low in 2021 (Fig. 2c). WGOM and GBK stocks contributed an approximately equal proportion of medium-sized cod to the regional abundance and followed similar abundance trends, while EGOM and SNE stocks slowly declined from the late 1980s to the late 1990s. SNE stock abundance has since remained low, but the EGOM stock has slightly increased since 2014. The regional COG has remained near the northwestern edge of the Georges Bank for the entire modeled period but has consistently shifted northward and eastward since approximately 2011 (Fig. 3c, i). Stock-specific COGs indicated that EGOM and GBK stocks have shifted northward and eastward over the modeled period (Fig. 4b). The WGOM stock has shifted consistently northward over the modeled period, but only began shifting consistently eastward in 2011. The SNE stock COG has shifted slightly south, but has no consistent east-west trend. The southwestern range edge movement matches that of the WGOM stock COGconsistent northward movement over the modeled period and eastward movement beginning in 2011, resulting in a northward displacement of nearly 75 km and eastward displacement of 30 km (Fig. 5c, i). The northeastern range edge has remained near the same east-west location but shifted approximately 100 km south from 1982 to 2011, matching a period of declining abundance in the EGOM stock. Since 2011, the northeastern range edge has rapidly returned to near its time-series northern extent. Range compressions and expansions matched trends in range edge shifts. The distance between the north and south range edges was consistent from 1982 to 1990, compressed by nearly 80 km from 1990 to 2010 due to declines of edge populations in the SNE and EGOM stocks, and regained approximately 42 km from 2010 to 2021. The distance between the east and west range edges did not fluctuate as much; it varied little from 1982 to 2011 but compressed by approximately 29 km from 2011 to 2021.

Model outputs for the fall time series resulted in similar patterns of spatial density as that of the spring time series (Fig. S7b). The index of relative abundance was lower in the fall than in the spring and has generally declined since the early 1990s, with the time-series achieving its lowest abundance in 2021 (Fig. 2d). WGOM and GBK stocks have contributed the most to the regional abundance over the time series. Abundance of the WGOM stock was consistent until 2018, at which point it began to decline. Abundance of the GBK stock has declined since the late 1990s. The EGOM stock began to decline in the late 1980s, with only a slight rebound beginning in the mid-2010s. The regional COG has shifted northward from the northwestern edge of Georges Bank into the WGOM stock area since approximately 2010 (Fig. 3d, j). Fall stock-specific COGs were similar to spring stock-specific COGs, both in location and directional trends (Fig. 3e). Fall range edges were also similar to spring range edges (Fig. 5d, j). Southwestern range edges remained at the same east-west location, but shifted more than 110 km north over the modeled period. Northeastern range edges shifted consistently south until 2011. The slight recovery of the EGOM stock was likely the driving force behind a return of the northeastern range edge to near its time-series northern maxima by 2021. These range shifts resulted in a north-south range compression of 98 km between 1982 and 2011, with a recovery of only 62 km between 2011 and 2021. East-west range compression was less severe, but there is evidence of compression intensifying between 2011 and 2020.

### 3.3 Large cod

Results from the drop-one density covariate significance testing indicated that the predicted likelihood of cobble, sand, and mud were not significant density covariates for large cod, and so they were excluded from all large cod model runs. In the spring time series, large cod were present at highest densities at the Northeast Peak region of Georges Bank and in the western Gulf of Maine around the edges of Wilkinson Basin on Stellwagen Bank, Jeffreys Ledge, and Cashes Ledge (Fig. S8a). The index of relative abundance peaked in 2003 and declined to a time-series low by 2021 (Fig. 2e). The GBK stock has historically contributed the most to large cod regional abundance, followed by the WGOM stock, and the stocks have similar trends in abundance over time. The regional COG was typically in the area of Georges Shoal, halfway between the regions of high density. Regional COG has moved consistently north and east since the early 1990s (Fig. 3e, k). Both the WGOM and GBK stock-specific COGs have moved steadily northward over the time series, but only GBK stock COG has moved consistently eastward (Fig. 4c). WGOM stock COG moved steadily west from 1982 until 2010, at which point it began moving rapidly east and surpassed its time-series eastern maximum in 2018. The spring southwestern range edge has shifted nearly 103 km northward over the time series and nearly 60 km eastward since 2011 (Fig. 5e, k). The northeastern range edge remained stationary in an east-west direction but shifted approximately 69 km south over the time series. These shifts resulted in a severe habitat compression; distance between the north and south range edges decreased by 171 km over the time series, and distance between the east and west range edges decreased by 133 km.

The density patterns of large cod in fall were similar to those in spring, but with lower density south of Cape Cod and in Georges Shoals (Fig. S8b). Regional abundance was lower in the fall than in the spring, with a peak abundance in 1989 and time-series low abundance in 2021 (Fig. 2f). Trends in abundance were strongly influenced by the GBK and WGOM stocks. GBK stock abundance increased through the early 2000s, after which point it began to decline. Interannual variability of WGOM stock abundance was high but did not decline at the same rate as the GBK stock, which increased its influence on large cod spatial dynamics in the later years of the time series. Like the spring regional COG, fall regional COG has moved consistently north since the early 1990s (Fig. 3f, l). But unlike the spring results, fall regional COG has shifted west, likely reflecting proportionally lower GBK stock contribution to the population. Trends in fall stock-specific COG movement were similar to spring stock-specific COG movements (Fig. 4f). Fall range edges indicated a rapid northward population shift and compression; the southwestern range edge moved over 100 km north and the northeastern range edge moved 38 km south over the time series (Fig. 5f, l). Range edges did not move in a consistent east-west direction over the time series. This resulted in a 130 km north-south habitat compression, with some evidence of east-west compression beginning in 2016.

### 3.4 Habitat associations

Depth, bottom water temperature, and the probability of encountering gravel were the most influential habitat covariates for all three size classes of cod for both linear predictors (Fig. 6, Fig. 7). All size classes had a clear unimodal relationship between depth and presence rate (first linear predictor), with an optimum depth between 40 and 65 meters and a monotonically decreasing presence rate past the optimum (Fig. 6b, j, s). The relationship between depth and positive catch rate (second linear predictor) was inconsistent across size classes (Fig. 7b, j, s). Small cod positive catch rate was highest in extreme shallows and decreased rapidly until approximately 60m, after which the rate of decrease was slower. Medium cod positive catch rate was also highest in extreme shallows and decreased until approximately 320m, after which it slowly increased. It should be noted that samples are sparse in waters deeper than 320m (<0.08% of all data), so these results may not be accurate. Large cod positive catch rates did not fluctuate much with depth, though the optimum was at approximately 45m.

**Figure 6:**
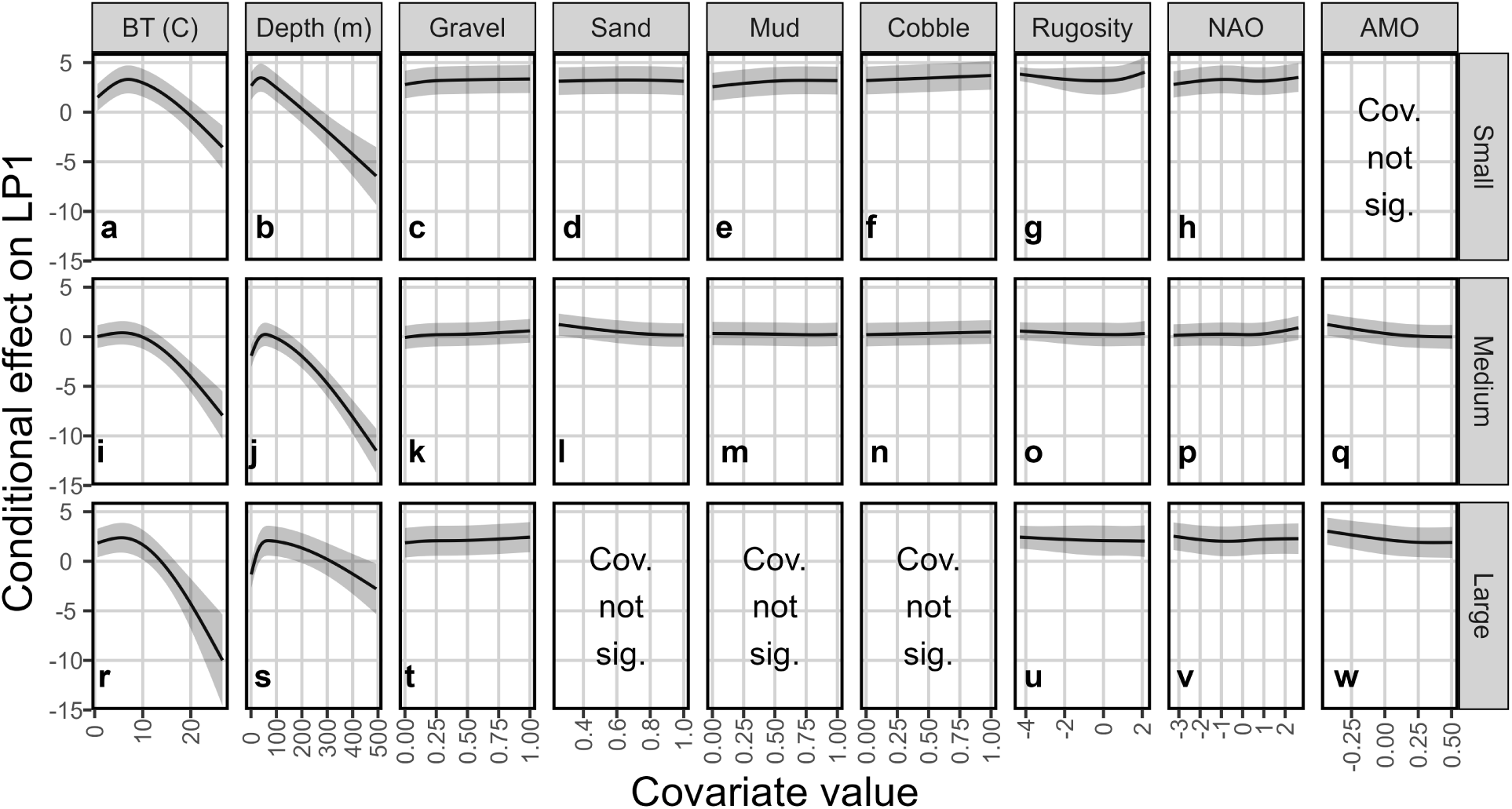
Conditional effects for density covariates on the first linear predictor (presence/ absence) within the a) small cod, b) medium cod, and c) large cod models. Conditional effects report the partial effects of each density covariate to linear predictors when all other density covariates are held fixed..

**Figure 7:**
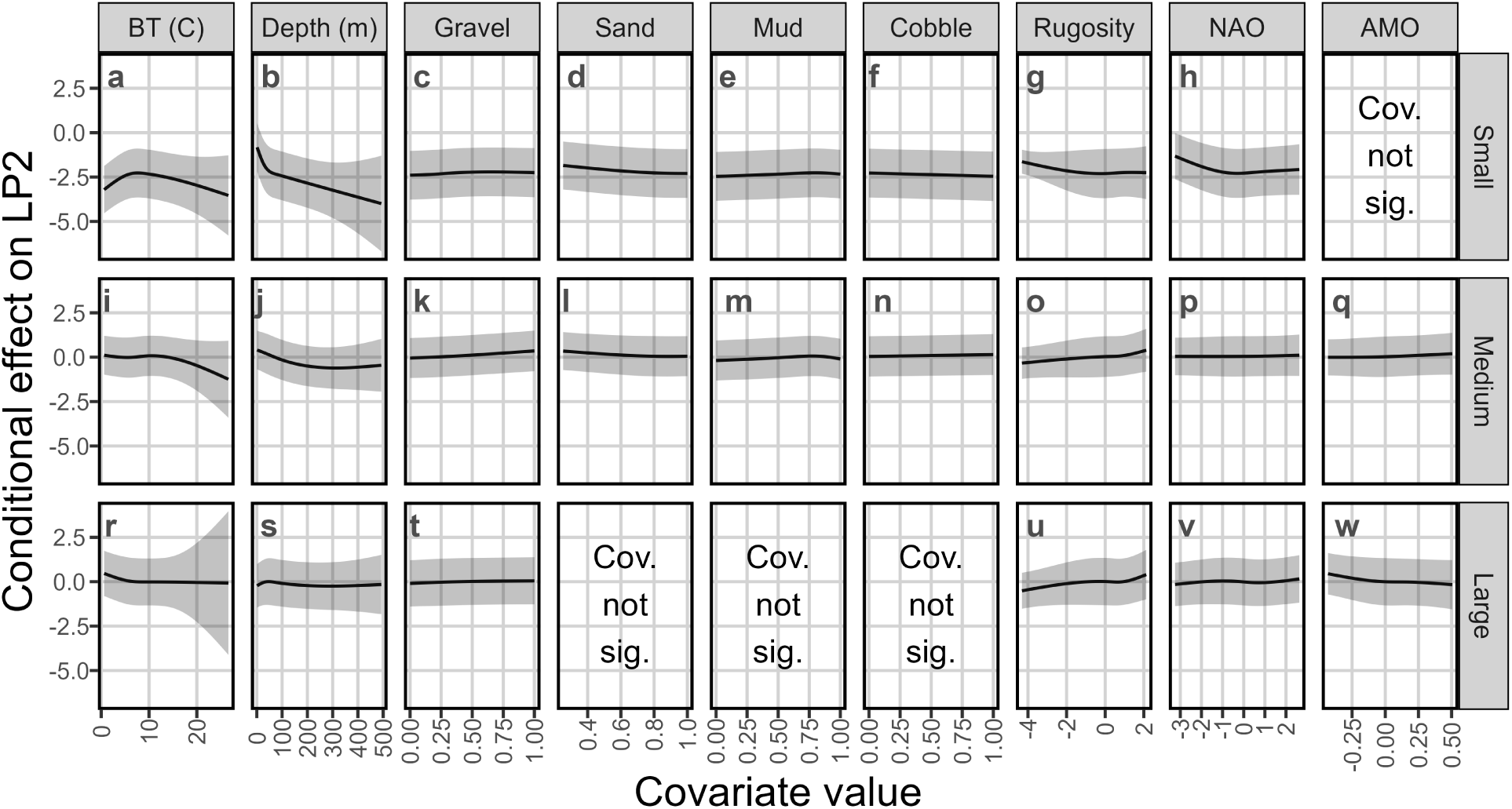
Conditional effects for density covariates on the second linear predictor (positive catch rate) within the a) small cod, b) medium cod, and c) large cod models. Conditional effects report the partial effects of each density covariate to linear predictors when all other density covariates are held fixed.

All size classes also had a clear unimodal relationship between bottom water temperature and presence rate, with optimum bottom temperature between 5.6 and 6.9řC and monotonically decreasing presence rate with warming bottom temperatures (Fig. 6a, i, r). There was a unimodal relationship between bottom temperature and positive catch rate for small cod, with optimum bottom temperature at approximately 7.7řC (Fig. 7a). Medium cod had a bimodal relationship between bottom temperature and positive catch rate, with local maxima at both 1řC and 10řC (Fig. 7i). Large cod positive catch rate was highest at approximately 1řC and decreased with increasing bottom temperatures (Fig. 7r). Few samples were taken in areas and times where bottom water temperature was below 2řC (<0.12% of all data), so the relationship between linear predictors and temperature may not be accurate at these extreme cold temperatures.

The probability of encountering gravel had a monotonically positive relationship with presence rate across all size classes (Fig. 6c, k, t) and a monotonically positive relationship with positive catch rate for medium and large size classes (Fig. 7k, t). For small cod, areas with a probability of encountering gravel beyond 65% had a slight decrease in positive catch rate (Fig. 7c). The remaining habitat covariates had limited influence on the two linear predictors and generally inconsistent results across size classes.

### 3.5 Unidentified drivers of spatio-temporal effects

Seasonal regional COGs for all size classes were compared with and without Gaussian Markov Random Fields (Fig. S9). Models for the small size class showed little difference in COGs between the two model types, implying that most major drivers of small cod COG variability are likely explicitly included in the models (Fig. S9a-b, g-h). The medium size class models consistently predicted COGs an average of approximately 20 km further east with GMRF turned on (Fig. S9i-j). There was also a temporal trend in the difference in northings between the GMRF On and Off models for this size class; in the first 20 years of the time series (1982-2001), GMRF On models predicted COG an average of 12-13 km farther north than the GMRF Off models. However, in the latter 20 years (2002-2021), this was reduced to only 2-3 km difference (Fig. S9c-d). These results suggested that much of the variability in COG for this size class was driven by sources not explicitly identified in the model, and that the unidentified driver may have a temporally variable influence that decreased over time. For the large size class, both GMRF On and GMRF Off models consistently showed a similar northward shift in regional COG (Fig. S9e-f). However, eastings were not as consistent between models. This error also had a temporal trend, with smaller differences in the first 20 years of the time series than in the latter 20 years (Fig. S9k-l). This finding was particularly evident in the fall time series. Again, this indicated that the model does not explicitly incorporate at least one major driver of large cod seasonal COG variability, and that this unidentified driver may have temporally variable influence.

## 4 Discussion

Atlantic cod stock assessments rely heavily on design-based indices of abundance built on data gathered with bottom trawl surveys. Despite evidence of higher cod abundance in areas with complex bottom habitats, locations with large-grain sediments or high rugosity are avoided or sampled at lower frequency with bottom trawls due to the risks of damaging equipment. This is concerning to fishing industry stakeholders, who believe that limiting information from complex habitats will not accurately reflect cod spatial distribution or abundance, therefore reducing precision and accuracy of design-based indices of abundance estimated from available data (Grabowski et al. 2020). Indeed, common concerns regarding design-based indices of abundance include mismatches between the spatial distribution of the stock and the spatial footprint of data collection, insufficient within-strata homogeneity for environmental conditions or species density, and inability to estimate density in un-or under-sampled regions (Thorson et al. 2015a; Adams et al. 2021; Cacciapaglia et al. 2024). Geostatistical models are gaining popularity in stock assessment methods because they can account for spatial autocorrelation and estimate smoothed surfaces of population density, resulting in more precise indices of abundance (Thorson et al. 2015a). However, the scale of spatiotemporal delta-models models like VAST, or their ability to estimate absolute abundance rather than relative or proportional abundance, is highly sensitive to model specifications (Thorson et al. 2021). This study was focused on the relative abundance of the biological stock areas and possible shifts in distribution rather than estimating absolute abundance. Therefore, we caution against interpreting our model-based indices as representative of absolute abundance. Future work could include both comparing our model-based indices to design-based indices and a model selection procedure to best estimate absolute abundance.

VAST models provide a flexible but robust framework to integrate data across multiple surveys and better assess the relative abundance of groundfish stocks across all habitat types within their spatial ranges. Here, VAST has allowed for the estimation of spatial, temporal, and spatiotemporal correlation of cod abundance, as well as the inclusion of vessel effects, to facilitate the combination of observation data from multiple sources with varied protocols. The model of cod spatial density created in this study utilized observation data from a suite of gear types and survey platforms, which bridged some of the data gaps inherent to models built with only bottom trawl survey data. Density covariates were used to interpolate cod spatial density in un-or under-sampled areas. The quality of model outputs is, in part, reliant on the quality of environmental and habitat data as density covariates. Though we used the best available data for density covariates, there is room for improvement in the confidence of underlying nearshore bottom water temperature data and sediment distribution data. Regardless, this integrated modeling process and its resulting indices of relative abundance may be better regarded by industry stakeholders than models built with only bottom trawl observation data.

VAST results highlighted the persistence of small patches with relatively high spatial density. Density values within these patches varied with abundance but remained elevated as compared to the rest of the spatial domain. This is consistent with the basin model of geographic distribution theory, in which density-dependent habitat selection leads to distribution contraction into optimal habitats (as predicted by the optimal foraging theory and the ideal free distribution model, Fretwell and Lucas (1969)) when populations are in decline (MacCall 1990). Atlantic cod stock distributions across regions, size ranges, and time periods have been identified as likely following density-dependent habitat selection processes, especially in periods of population decline (Swain and Wade 1993; Rose and Kulka 1999; Blanchard et al. 2005; Tamdrari et al. 2010; Pershing et al. 2015; Thorson et al. 2016; Li et al. 2018). It is likely that direct environmental forcing or behavioral mechanisms that support density-dependent habitat selection have condensed cod to spatiotemporally variable optimal habitat patches, and areas outside these patches are now less suitable for sustained occupation.

As abundance declines, density within these patches decreases at a much slower rate than in other areas, resulting in greater proportions of the overall cod regional abundance contained within these relatively small successful patches. The continued success of these patches despite declining overall regional abundance is further evidence for the basin model, and hints at a source-sink dynamic where high-density patches are likely natal areas for cod that range more widely in the study area (Kritzer and Cadrin 2012). Genetic research has already identified many of these patches as important spawning grounds, which has informed the new stock structure utilized in management (Kovach et al. 2010; Zemeckis et al. 2014b; Clucas et al. 2019; McBride and Smedbol 2022). Further analysis could determine the degree to which environmental forcing (through spatiotemporally dynamic conditions like bottom water temperatures) and intraspecific interactions shape cod distributions within each stock area.

Range shifts towards more northerly and deeper waters have been observed in many North Atlantic marine species (Perry et al. 2005; Nye et al. 2009; Pinsky et al. 2013; Fredston et al. 2021). These climate-mediated shifts are associated with range expansions and increases in abundance for some northwest Atlantic fishes like summer flounder (Nye et al. 2009) and Georges Bank stocks of haddock (Wang et al. 2024). For cod, model results generally indicate a northward and offshore shift in center of gravity, a north-south range contraction, and a decline in abundance over the time series. These changes in spatial dynamics are occurring simultaneously with temperature-linked reductions in reproduction and growth (Planque and Frédou 1999; Drinkwater 2005; Fogarty et al. 2008; Pershing et al. 2015), which emphasizes the role of habitat preferences for colder waters in contributing to range contractions and decreasing abundance, particularly in the inshore and southern areas of cods spatial range.

Intense surface and bottom water warming signals within the Northeast US continental shelf 2008-2011 may have signaled a regime shift into warmer waters (Friedland et al. 2020). Models without fine-scale interpolation (through GMRFs) were also less able to track rapid COG shifts beginning in this period. As the northwest Atlantic, and specifically the Gulf of Maine-Georges Bank ecoregion, warm at a rapid rate (Pershing et al. 2015), it should be expected that the spatial dynamics of cod will change rapidly. Thus, it is imperative that climate signals be included in stock assessment methods. Model outputs indicate that a consequence of warming waters will be a condensed spatial distribution of cod into increasingly smaller patches of suitable habitat. Successful management efforts will need to consider the potential negative effects of reduced patch size and overall suitability of the Northeast US continental shelf under warming conditions to cod recruitment and survival. Environmental forcing may strongly affect the small size class, as its distribution is concentrated within shallow nearshore waters most vulnerable to increasing bottom water temperatures (Kavanaugh et al. 2017). The benthic characteristics and temperatures of high-density patches of small cod should be of particular concern to managers, as juvenile mortality rates have been hypothesized as more important to cod population growth than adult survival and fecundity (Wright 2014).

Several areas closed to groundfishing are among the persistent high-density patches. For example, large cod were consistently present at high densities in the Western Gulf of Maine Closed Area. Small cod were consistently present at high densities in nearshore waters within and around the Great South Channel Habitat Management Area. The continued decline of the regional abundance despite the creation of areas closed to fishing and increased fishing regulations points to a multitude of compounding problems beyond fishing alone. However, previous research has noted that even slight fishing pressure compounded with abiotic stressors will extend the rebuilding process (Pershing et al. 2015), and that fishing pressure may have increased selection for sedentary fish unlikely to colonize suitable habitats outside of closed areas (Sherwood and Grabowski 2010).

The application of a spatiotemporal delta-model, like VAST, provides the ability to assess the proportion of spatial density variation attributable to specific habitat features. The modeled relative density of all cod size classes in this study indicated strong preferences for depth and bottom temperature, and weaker preferences for substrate gravel content. As explained earlier in this section, we cannot assert that our models are scaled appropriately to calculate absolute abundance. However, the size-specific habitat preferences, spatially-explicit high-density patches, and relative abundance between biological stock areas resulting from this study could be used to inform novel survey designs that would increase accuracy and precision of design-based indices of abundance. Our results support shifting from depth-and latitude-based strata to dynamic stratification, in which sampling strata are built on multiple important habitat characteristics and the likelihood of encountering cod. Dynamic stratification has been previously suggested by industry members and Grabowski et al. (2020). The complicated interplay of cod population dynamics, stock structure, habitat preferences, and a dynamic environment make assessing the status of cod stocks and enacting effective management very challenging. The results of this study are clear evidence that the spatial dynamics and habitat preferences of biological stocks and age classes are variable, and this must be accounted for in stock assessment and when proposing measures to rebuild the population.

## Acknowledgements

The authors would like to thank vessel crew, research staff, and survey managers at Fisheries and Oceans Canada, NOAA Northeast Fisheries Science Center, Atlantic States Marine Fisheries Commission, UMass Dartmouth School for Marine Science & Technology, Massachusetts Department of Marine Fisheries, University of Rhode Island Graduate School of Oceanography, Rhode Island Department of Environmental Management, New Hampshire Fish and Game, Maine Department of Marine Resources, Maine Center for Coastal Fisheries for conducting the surveys used in this study. Further, the authors thank Felipe Restrepo, Brad Harris, and Hubert du Pontavice for sharing envioronmental data used as density covariates. Finally, the authors thank Jackie Odell, Vito Giacalone, Andrew Allyn, and Alex Hansell for advice and comments during the data processing and analysis stages.

## Author contributions

Conceptualization: GS, LK, JG Data curation: KL, LK Formal analysis: KL Funding acquisition: GS, LK Methodology: KL Resources: KL, LK Supervision: GS, LK Visualization: KL Writing - original draft: KL Writing - review & editing: KL, JG, GS, LK

## Data availability

All code and most data are available on request to the corresponding author. Raw results from trawl surveys and sediment probaility rasters cannot be shared, as per data sharing agreements.

