## Supplemental Text for "Spatial density and habitat associations of Atlantic Cod on the Northeastern US Continental Shelf"

**1	Density Covariates**

**1.1	Other covariates investigated for inclusion**

Sea surface temperature (SST) could affect spawning and recruitment success, and so was included as a potential density covariate (Planque and Frédou 1999; Drinkwater 2005; Fogarty et al. 2008; Pershing et al. 2015; Klein et al. 2017). Though most surveys measured and reported SST for every observation, the empirical dataset includes many missing SST values. VAST cannot tolerate missing values in density covariates and removing a large proportion of data was undesirable. Therefore, NOAA’s 1/4° spatial resolution daily Optimum Interpolation SST (OISST) data product was used to fill gaps. OISST daily rasters were pulled from NOAA data sources and SST was extracted at observation locations. OISST values were compared to field measurements, when available, and the two were found to be generally similar.

**1.2	Collinearity tests**

Before inclusion in model selection, potential density covariates were tested for collinearity. High correlations were found between two pairs: cobble and rock sediment probability, and OISST and bottom temperature. For the first collinear pair, rock sediment probability was removed, and cobble sediment probability was retained in the model. Grab and coring sampling methods (the bulk of sediment samples that support the sediment model) are unlikely to sample large-grain sediments like boulders effectively, and so the lower-quality data that feeds this model will inevitably result in a lower-quality and less reliable model of large-grain sediment distribution (Bachman et al. 2011). For the second pair of collinear variables, OISST was removed as it was expected that the distribution of groundfish like cod would be more directly affected by bottom temperature than sea surface temperature. Once these two collinear relationships were addressed, the remainder of density covariates were not correlated and therefore were tested for inclusion in the final model.

**2	Relative influence of each survey to indices of abundance**

**2.1	Methods**

The relative influence of each survey on model-based indices of abundance was of interest, given the issues the final models likely have with scalability to absolute abundance. To assess relative influence, a simple leave-one-out approach was used. For each size class, the final model structure was re-run 10 times, with a different survey excluded each time. Seasonal indices of relative abundance within each biological stock area were derived from the resulting spatiotemporal density models and compared to the index of relative abundance from the final model (with all surveys included). This was done to compare overall trends, identify any scaling issues, and to identify years in which the removal of a single survey resulted in large differences in estimated abundance.

Similarity of overall trends were assessed by treating all seasonal indices of relative abundance as time series with 40 steps (one step per year), then using the cross-correlation function (CCF) to complete pairwise comparisons of the time series from each of the drop-one-survey models to the final model. The null hypothesis of the CCF is that correlation between time series is not significantly different to 0 at the chosen lag value, or in other words, the time series are not correlated at that lag. It essentially calculates Pearson correlation coefficients for sets of a variable at different lag intervals. Approximate critical values at the α = 5% level (resulting in a 95% confidence interval around 0 correlation) for each lag can be calculated as ${\pm2}/\sqrt{n}$, where *n* is the length of the overlap between compared time series. As we are focused on whether compared time series move in the same direction at the same time, we are only interested in results at a lag of 0. We therefore have *n* = 40 time steps for each time series comparison, and the confidence thresholds are 0.31 and -0.31. If the correlation value is positive and exceeds the upper bounds of the calculated 95% confidence interval, there is evidence that the compared time series have similar trends. Correlation values between the 95% confidence interval bounds are evidence of no correlation between time series. Negative correlation values below the lower 95% confidence interval bound are evidence of significant negative correlation, or opposite trends. Cross-correlation values greater than 0.6 are typically considered to indicate strong relationships, and values greater than 0.8 indicate very strong relationships. We will report both significance and relative strength of time series cross-correlations.

Cross-correlation analysis is useful to assess the similarity of overall trends between indices of abundance but is not sensitive to scaling between time series. For example, the cross-correlation values of time series *x* and time series *x* * 2 or time series x + 100 would be 1, indicative of a perfect correlation. Further, it cannot identify specific years where the exclusion of a survey leads to large differences in estimated abundance. It is important for us to assess not just overall abundance trends, but also the magnitude of differences in estimated abundance across years. To identify differences in scaling, we compared the mean ratio of the abundance index from the drop-one models in a given year (*D_y_*) to the abundance index from the final model (*F_y_*). This mean ratio (*R*) is therefore calculated as

$$R = mean ({D_{y}}/{F_{y}})$$

and values closer to 1 indicate agreement in scale. This metric has been suggested by both Thorson et al. (2021) and Cacciapaglia et al. (2024) in the context of comparing VAST-based indices of abundance to a design-based index of abundance, but is also appropriate to assess scaling between two VAST-based indices.

Finally, we compared the 95% confidence interval of the final model index of relative abundance against the indices of relative abundance from the drop-one models. The number of years in which the drop-one indices (*D_y_*) fell outside the final model index (*F_y_*) were counted and called *TimeOut*. This metric was originally developed by Cacciapaglia et al. (2024) to compare similarity of design-based and VAST-based indices of abundance, but is also useful to identify differences between two VAST-based indices. If more than 5% of modeled time steps fell outside the 95% confidence interval, the excluded survey was identified as having high relative influence to that season and biological stock area. As each seasonal index of relative abundance has 40 time steps, the threshold is more than 2 instances of falling outside the confidence interval bounds. The calculated R and TimeOut values for each size – season – biological stock area index of relative abundance can be found in Table SM1.

**2.2	Results**

It should be expected that removing the NEFSC bottom trawl survey will significantly alter seasonal indices of relative abundance for all size classes and biological stock areas. It is the only source of offshore data across most of the modeled spatial domain and makes up 30% of total data volume. Indeed, seasonal indices of relative abundance of small and medium cod are below the 95% confidence bounds of the final model most of the time (>50% of time series) when this survey is not included in modeling efforts. Results from models with BTS data removed will not be included in the remainder of this section, as it would be unreasonable to remove this vital dataset from any model intended to realistically represent cod spatiotemporal density.

**2.2.1	Small cod**

For the small size class, the Massachusetts Department of Marine Fisheries inshore trawl survey (hereafter: MADMF inshore survey) was identified as influential to indices of abundance (Fig. SM1). This survey operates within the SNE and WGOM biological stock areas, covering most Massachusetts territorial marine waters. It has conducted spring and fall surveys since 1982. Cross-correlation analysis identified significant differences in WGOM – Spring abundance trends with the removal of the MADMF inshore trawl survey (cross-correlation: 0.20). The SNE – Spring and WGOM – Fall indices were significantly correlated with the final model indices in the same seasons and biological stock areas, but cross-correlation values fell below the value needed to be considered “very strong” relationships (cross-correlation values 0.65 and 0.79, respectively). Removing all data from this survey decreased estimated small cod abundance in all seasons and biological stock areas, and the notable 2003 abundance spike within the SNE and WGOM spring indices does not appear without these data. Estimated abundance is much lower in the SNE – Spring (R: 0.28, TimeOut: 40 years), WGOM – Spring (R: 0.39, TimeOut: 39 years), SNE – Fall (R: 0.74, TimeOut: 11 years), and WGOM – Fall (R: 0.69, TimeOut: 27 years) indices. Abundance is also underestimated for shorter periods in the EGOM and GBK spring indices. Clearly, VAST-based model indices without the MADMF inshore survey data are scaled much lower than models that include them.

Though CCF results did not indicate any other survey significantly altered abundance trends when removed, several were noted to have cross-correlation values beneath the threshold to have “very strong” relationships. In particular, the SNE – Spring indices generated with and without the Fisheries and Oceans Canada (DFO) trawl data had only a “moderate” relationship (cross-correlation value: 0.52). Removal of the DFO trawl data also led to periods of index scaling differences as indicated by the mean ratio and TimeOut metrics; comparison to the final model indicated long periods of overestimating abundance in the GBK – Fall index (R: 1.81, TimeOut: 26 years) and underestimating abundance in the SNE – Fall (R:0.79, TimeOut: 23 years) and GBK – Spring (R:0.98, TimeOut: 15 years) indices. It also led to shorter periods of underestimated abundance in the WGOM – Spring index and overestimating abundance in the EGOM – Fall and WGOM – Fall indices. Removal of the Rhode Island Department of Environmental Management (RIDEM) trawl led to underestimating abundance in the SNE – Spring index (R: 0.88, TimeOut: 14 years) and overestimating abundance for the fall index in the same area (R: 1.22, TimeOut: 4 years). Removal of the Maine-New Hampshire Inshore trawl survey generally underestimated abundance in the EGOM Spring (R:0.96, TimeOut: 12 years) and Fall (R:0.94, TimeOut:18 years) indices. Removal of the SMAST and MADMF Industry-based trawl surveys led to short periods (TimeOut: 3-6 years) of overestimated abundance for spring and fall WGOM indices and spring EGOM indices.

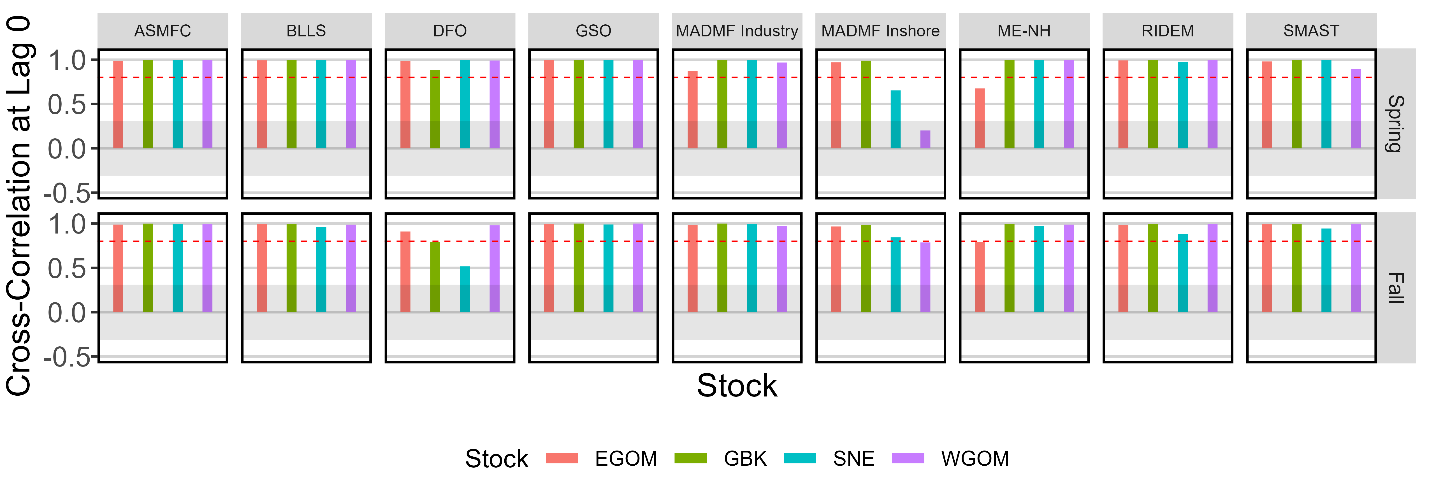

Figure SM 1: Cross-correlation function values at lag 0 comparing the final and drop-one seasonal indices of relative abundance within each biological stock area for the small size class. The shaded polygon indicates the region in which there is insignificant evidence to reject the null hypothesis of no correlation (α=0.05). The red dotted line is the threshold above which correlation can be considered “very strong” (0.8).

**2.2.2	Medium cod**

Cross-correlation analysis did not indicate that the removal of any survey would alter temporal trends in medium cod indices of relative abundance (Fig. SM2). However, GBK – Fall indices generated with and without the DFO trawl data only had “moderate” relationships (cross-correlation value: 0.59). All other index comparisons indicated very strong relationships (cross-correlation value > 0.8). Models without DFO trawl data generally estimated lower abundance, particularly in the GBK biological stock area. Interestingly, the GBK – Spring index without DFO trawl data changed from periodically estimating lower abundance 1988 – 2010 to estimating higher abundance 2011 – 2017. Removing the Maine-New Hampshire inshore trawl survey led to lower estimated abundance in the EGOM indices (Spring – R: 0.88, TimeOut: 7 years; Fall – R: 0.92, TimeOut = 5 years) and removing the NEFSC bottom longline survey led to lower estimated abundance in the WGOM indices (Spring – R: 0.84, TimeOut: 8 years; Fall – R: 0.81, TimeOut: 10 years) and the EGOM – Fall index (R: 0.74, TimeOut: 4 years). Removing the MADMF inshore trawl survey reduced expected abundance in the WGOM – Spring index (R: 0.87, TimeOut: 14 years) and increased expected abundance in the SNE indices (Spring – R: 1.16, TimeOut: 5 years; Fall – R: 1.53, TimeOut: 29 years). Removal of the Atlantic States Marine Fisheries Commission’s (ASMFC) northern shrimp trawl increased expected medium cod abundance in the EGOM and WGOM spring indices (Spring – R: 1.09, TimeOut: 4 years; Fall – R: 1.19, TimeOut: 19 years). Removal of the MADMF Industry-based trawl surveys led to a long period of overestimating cod abundance in the Spring – EGOM index (R: 1.33, TimeOut: 11 years), shorter periods of overestimating abundance in the Fall – EGOM and Spring – WGOM indices, and a few years of underestimating cod abundance in the Fall – WGOM index. Finally, removal of the RIDEM and SMAST surveys led to short periods (TimeOut: 3-5 years) of overestimated abundance in the fall SNE and spring WGOM indices, respectively.

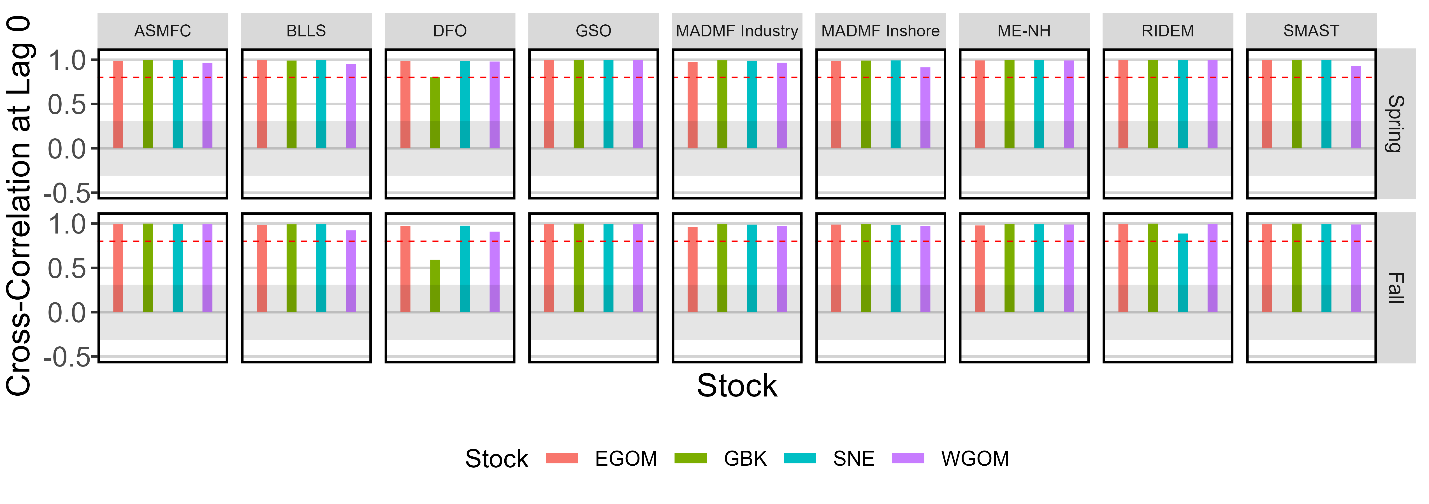

Figure SM 2: Cross-correlation function values at lag 0 comparing the final and drop-one seasonal indices of relative abundance within each biological stock area for the medium size class. The shaded polygon indicates the region in which there is insignificant evidence to reject the null hypothesis of no correlation (α=0.05). The red dotted line is the threshold above which correlation can be considered “very strong” (0.8).

**2.2.3	Large cod**

Large cod models were extremely sensitive to any data removal. Most drop-one models failed to complete, usually because they were unable to estimate spatial effects in the second linear predictor (L_omega2_z). Removing the DFO trawl survey data resulted in not only this issue but also a failure to estimate the temporal structure of spatiotemporal variation in the first linear predictor. Removal of the NEFSC BTS resulted in failure to estimate a gradient for geometric anisotropy for the second linear predictor. The only model that ran to completion was when the MADMF Inshore Trawl data were removed; this survey has a relatively small spatial footprint that mostly covers areas unlikely to be suitable for large cod, particularly in the fall time series.

**2.3	Discussion**

The rarity with which cross-correlation analyses indicated the removal of any survey (except the BTS) would significantly affect the trends of any seasonal, stock-specific index of cod relative abundance adds confidence to the interpretation of final VAST model results. A notable exception to this is the influence of the MADMF inshore survey on small cod spring indices of relative abundance within the WGOM biological stock area. Ecologically, it should be expected that the area covered by this survey within the Western Gulf of Maine (including Massachusetts Bay and Cape Cod Bay) would contain high densities of small cod, which prefer shallower and warmer waters than the larger size classes. There is little survey coverage of this area without this survey, so we are not recommending the removal of these data. Instead, this should be seen as a strong motivator to assess accuracy to absolute abundance via the model structuring decision tree suggested in Cacciapaglia et al. (2024) and comparison of design-based and model-based indices of relative abundance.
