## Supplementary figures and images for "Spatial density and habitat associations of Atlantic Cod on the Northeastern US Continental Shelf"

### Supplemental Figure 1

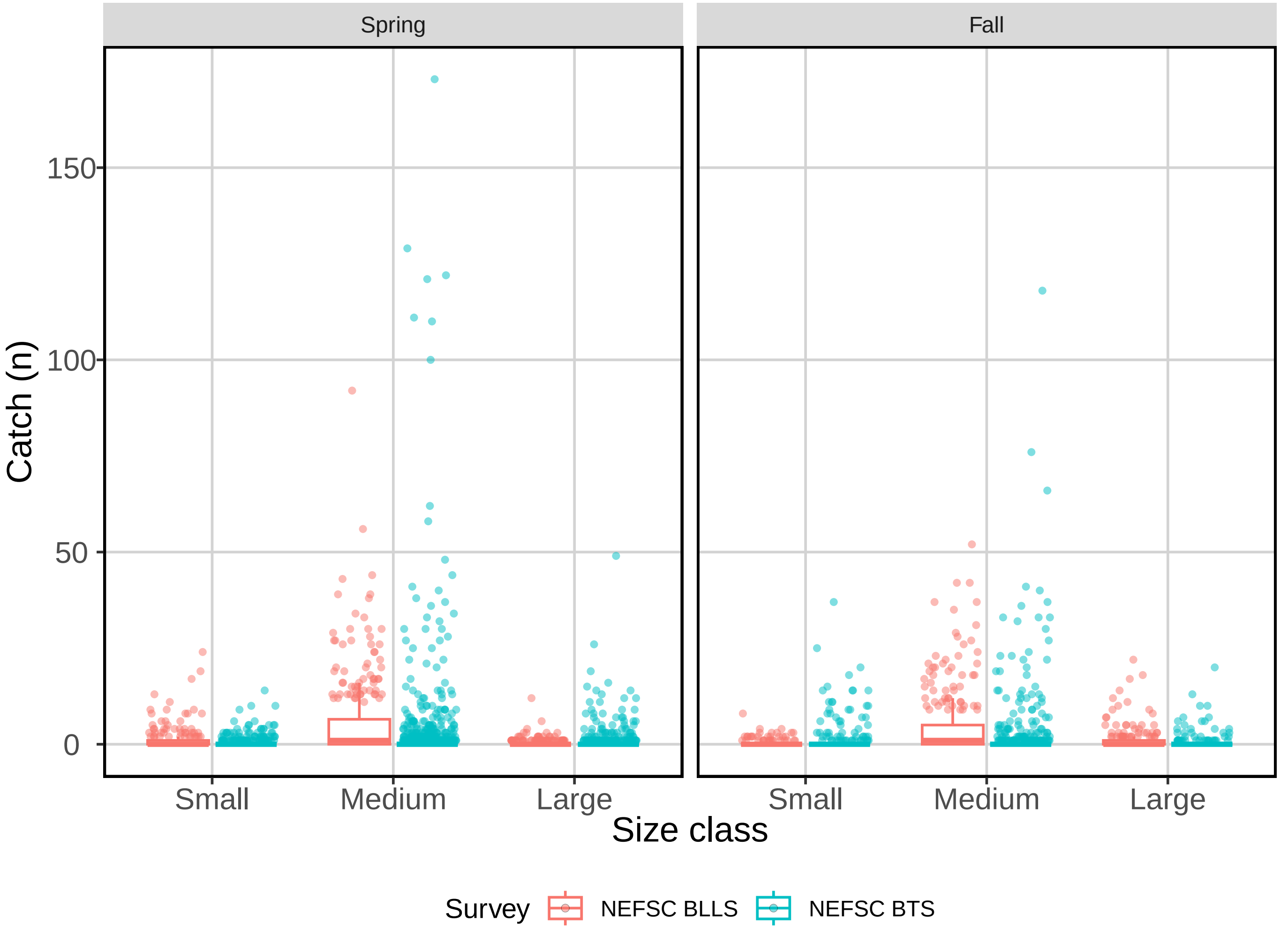

### Supplemental Figure 2a

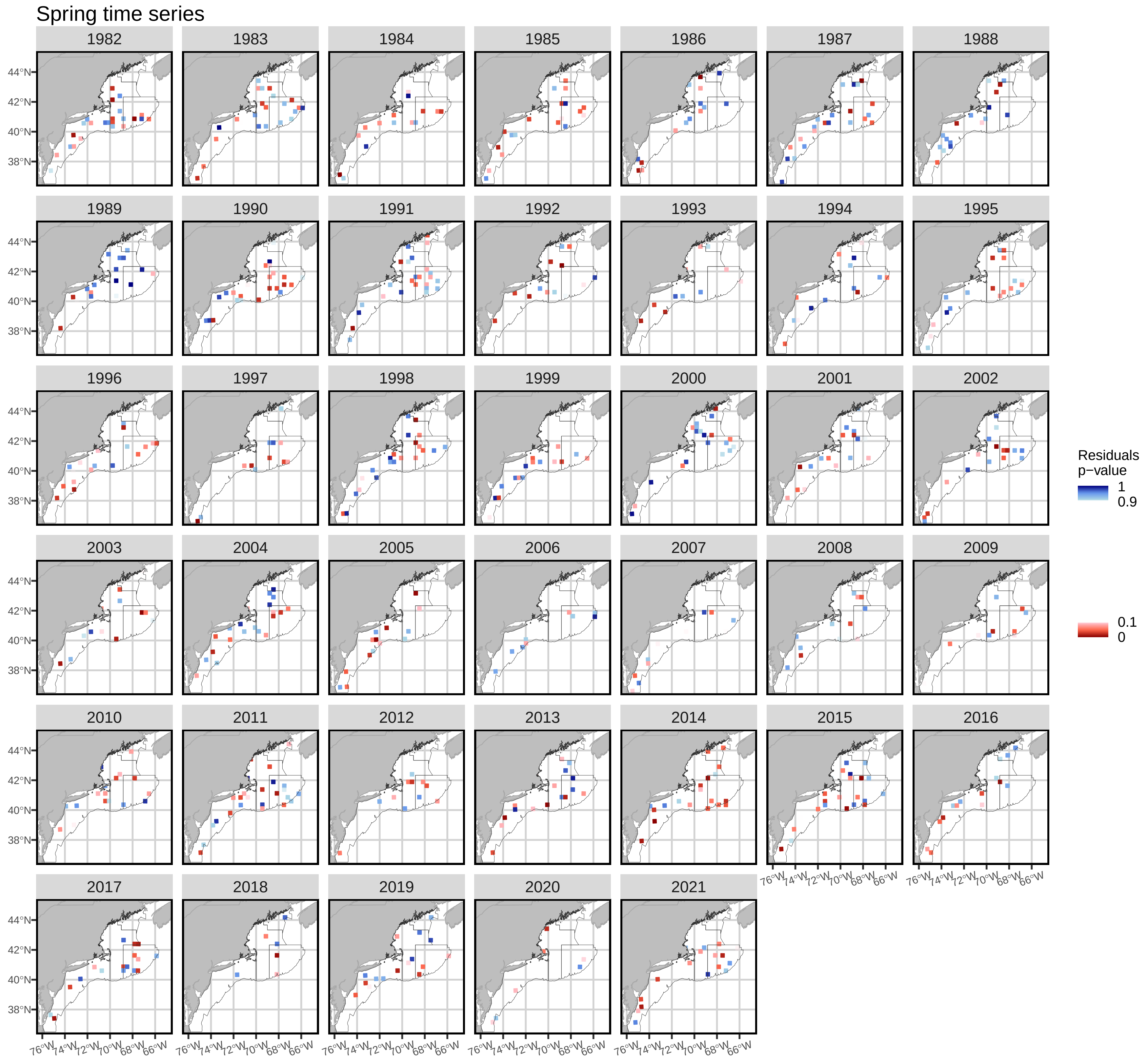

### Supplemental Figure 2b

##### Fall time series

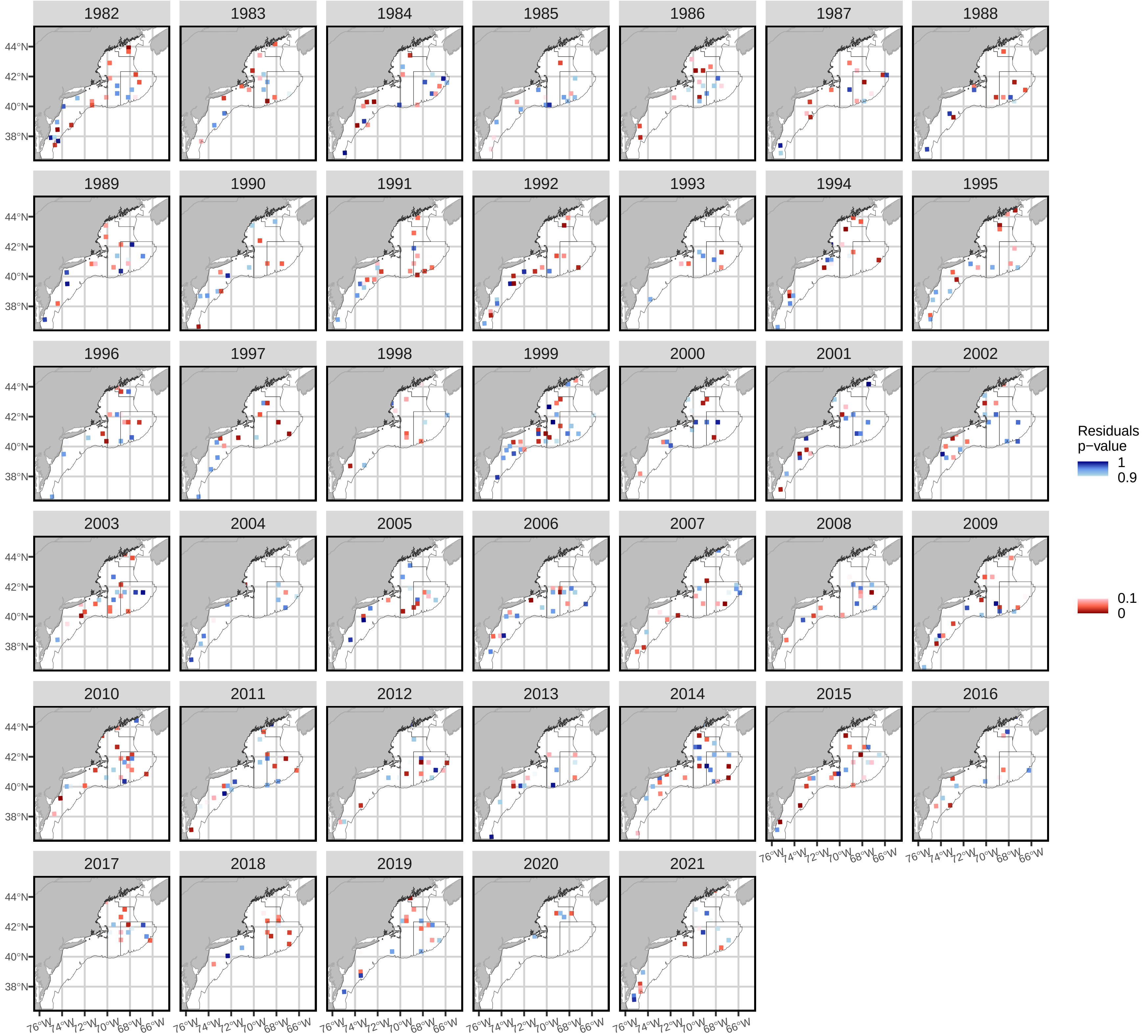

### Supplemental Figure 3a

##### Spring time series

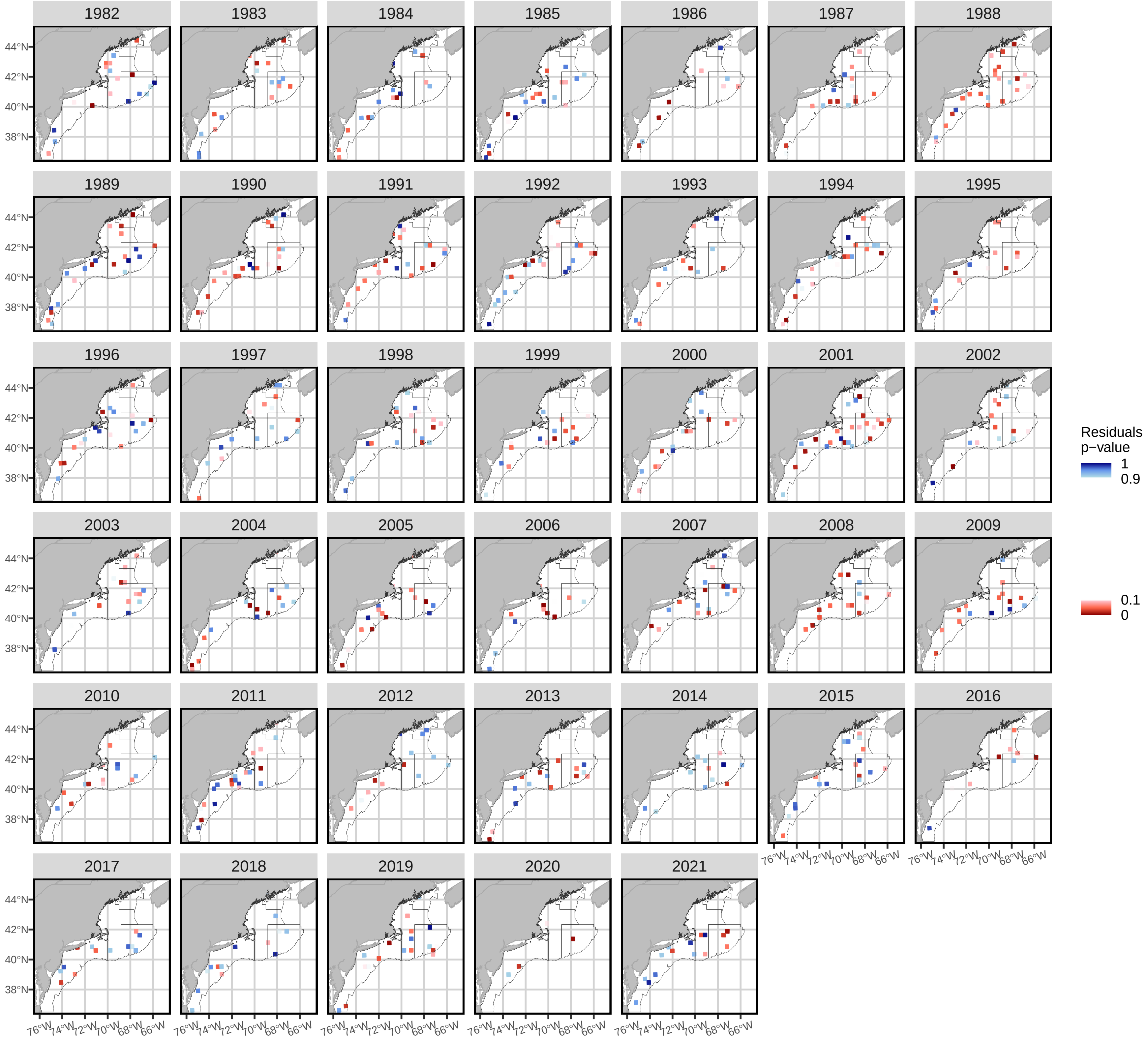

### Supplemental Figure 3b

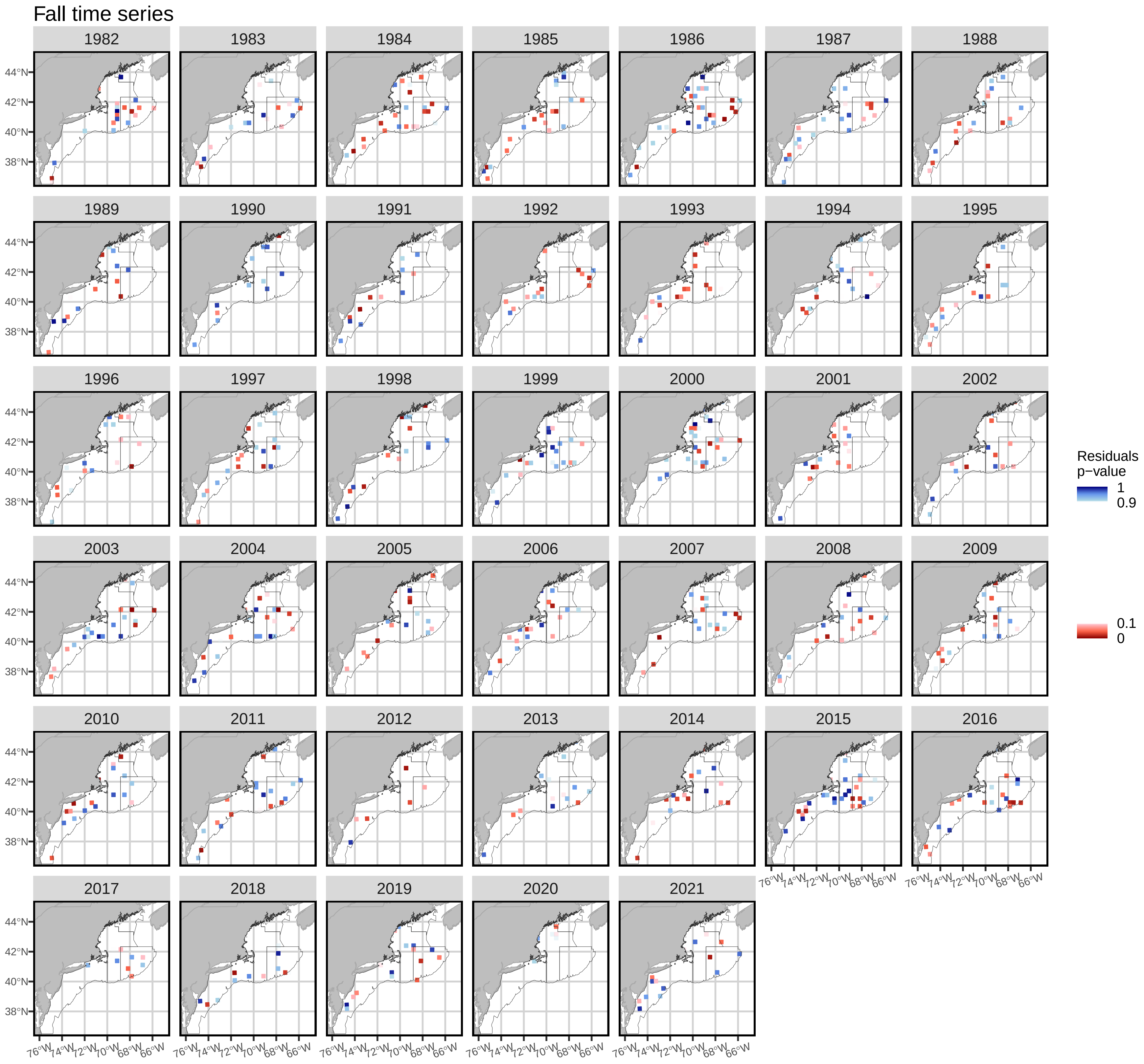

### Supplemental Figure 4a

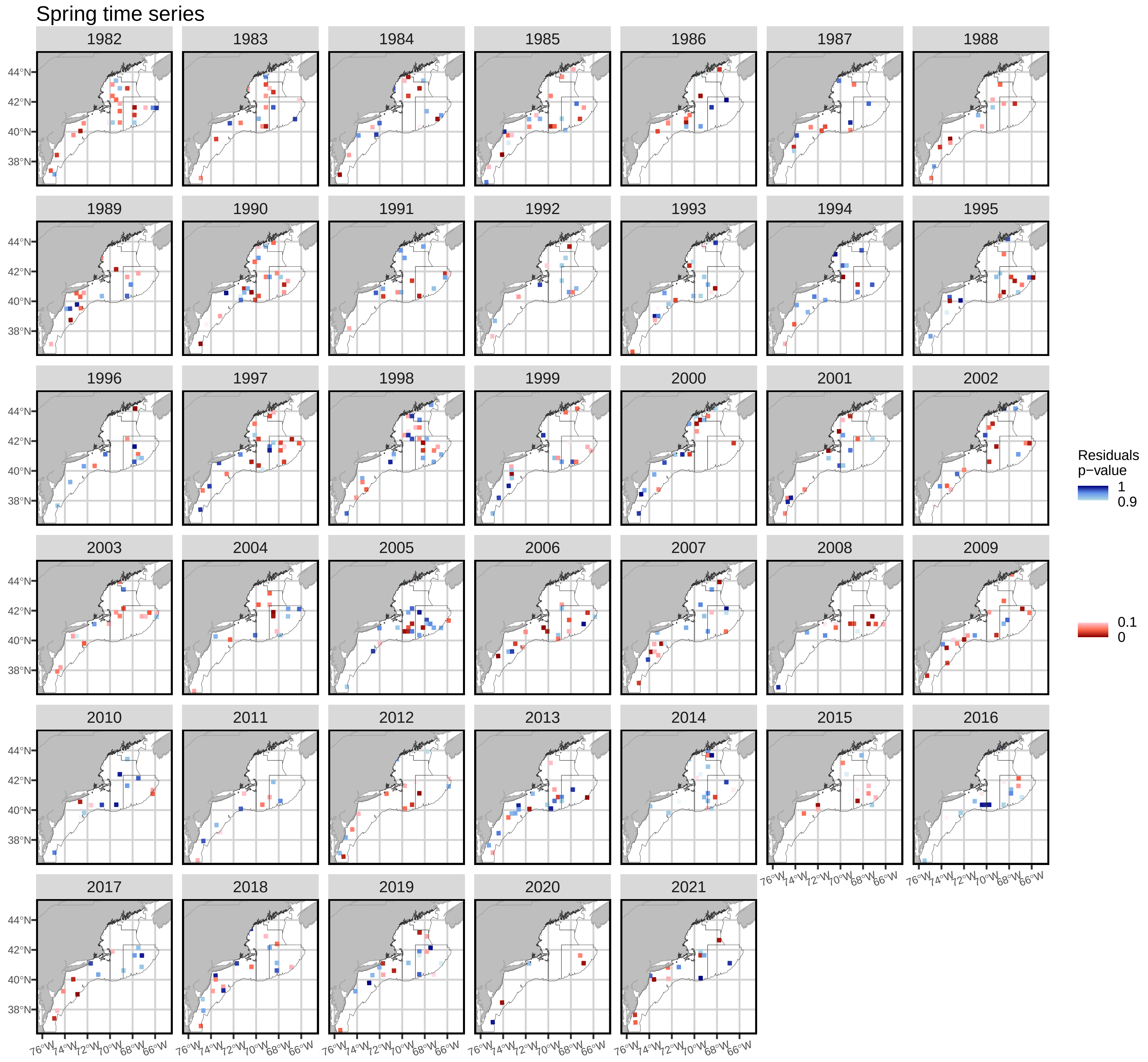

### Supplemental Figure 4b

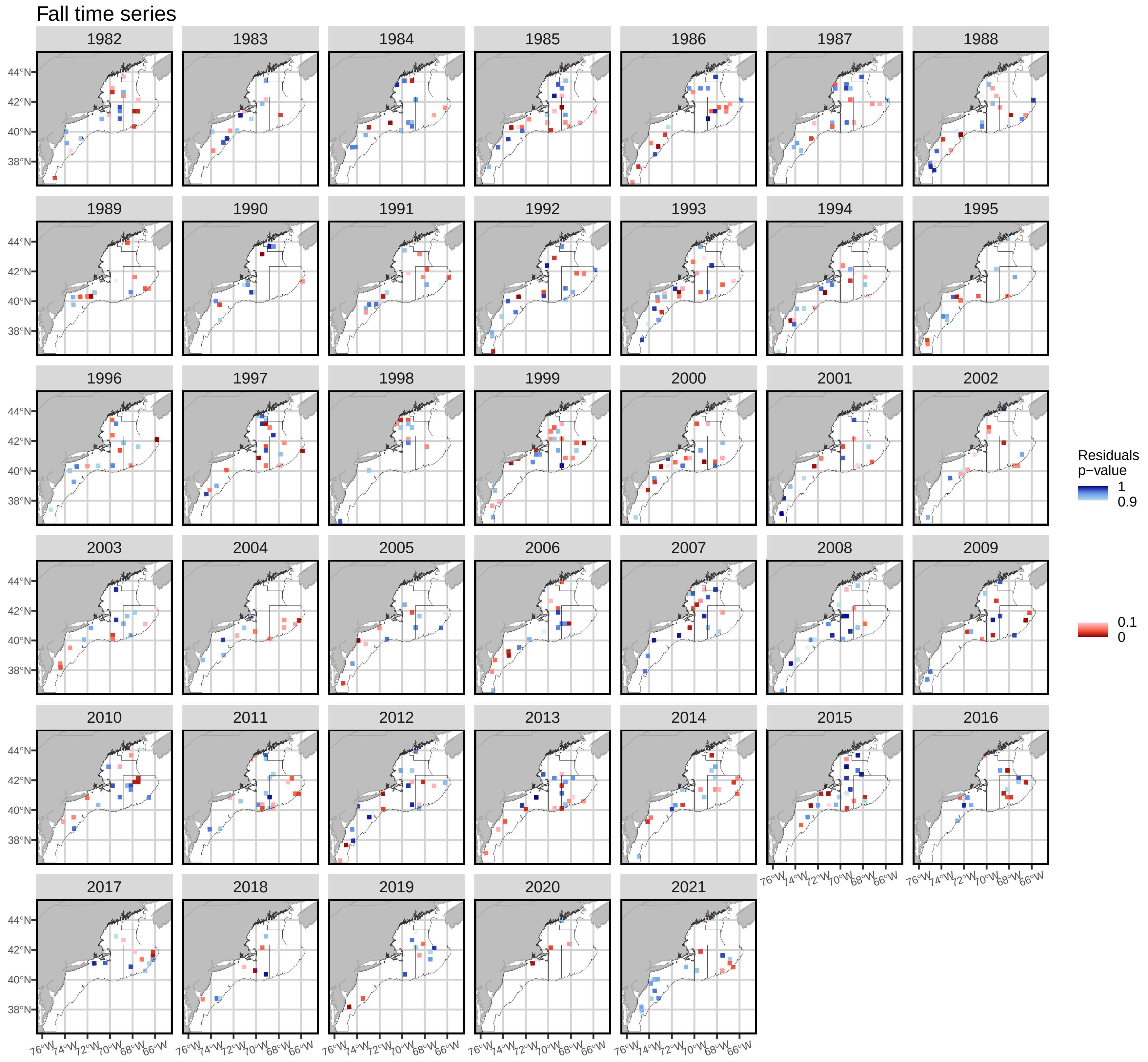

### Supplemental Figure 5

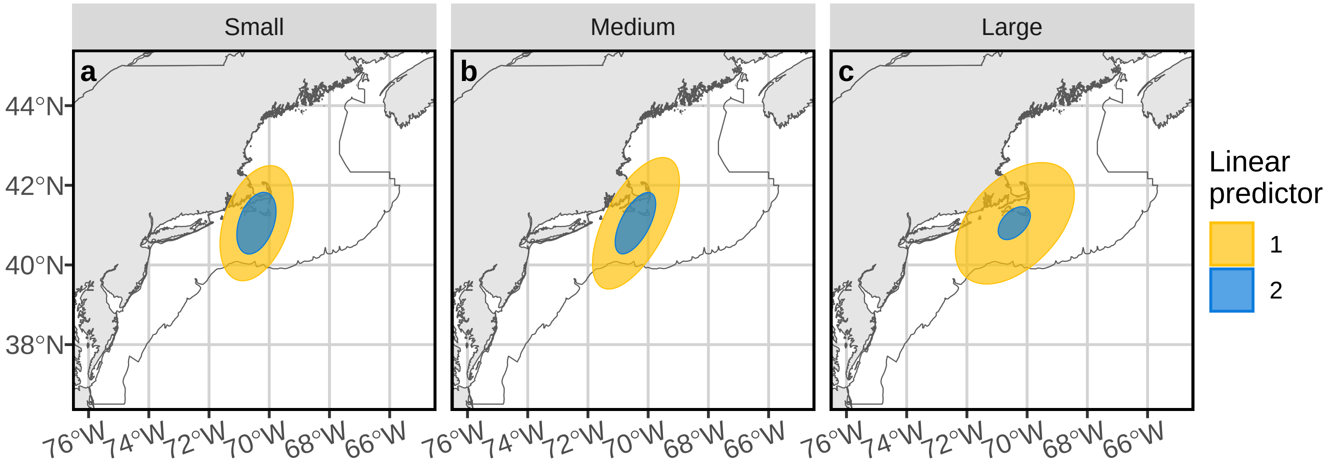

### Supplemental Figure 6a

#### Spring spatial density

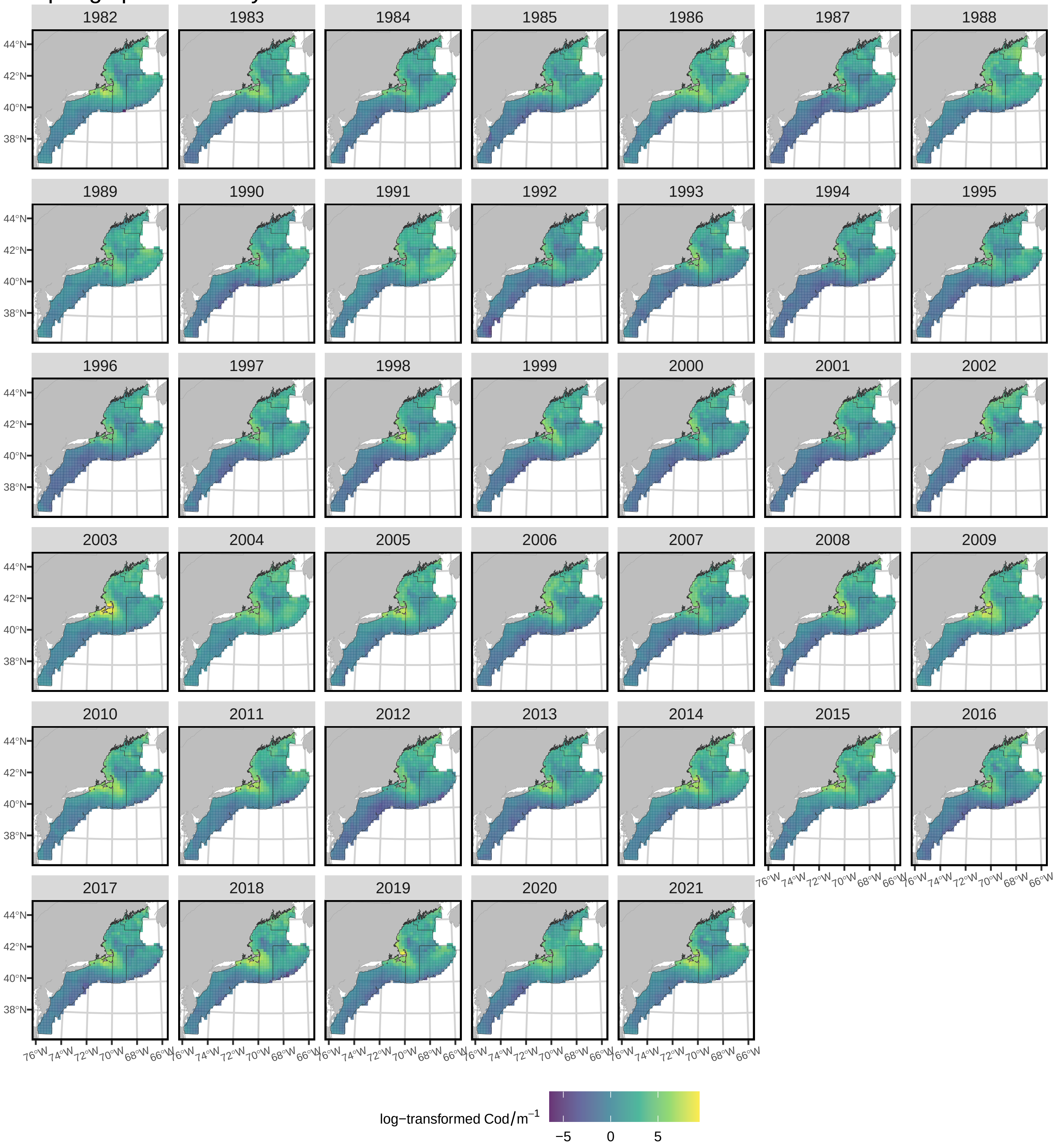

### Supplemental Figure 6b

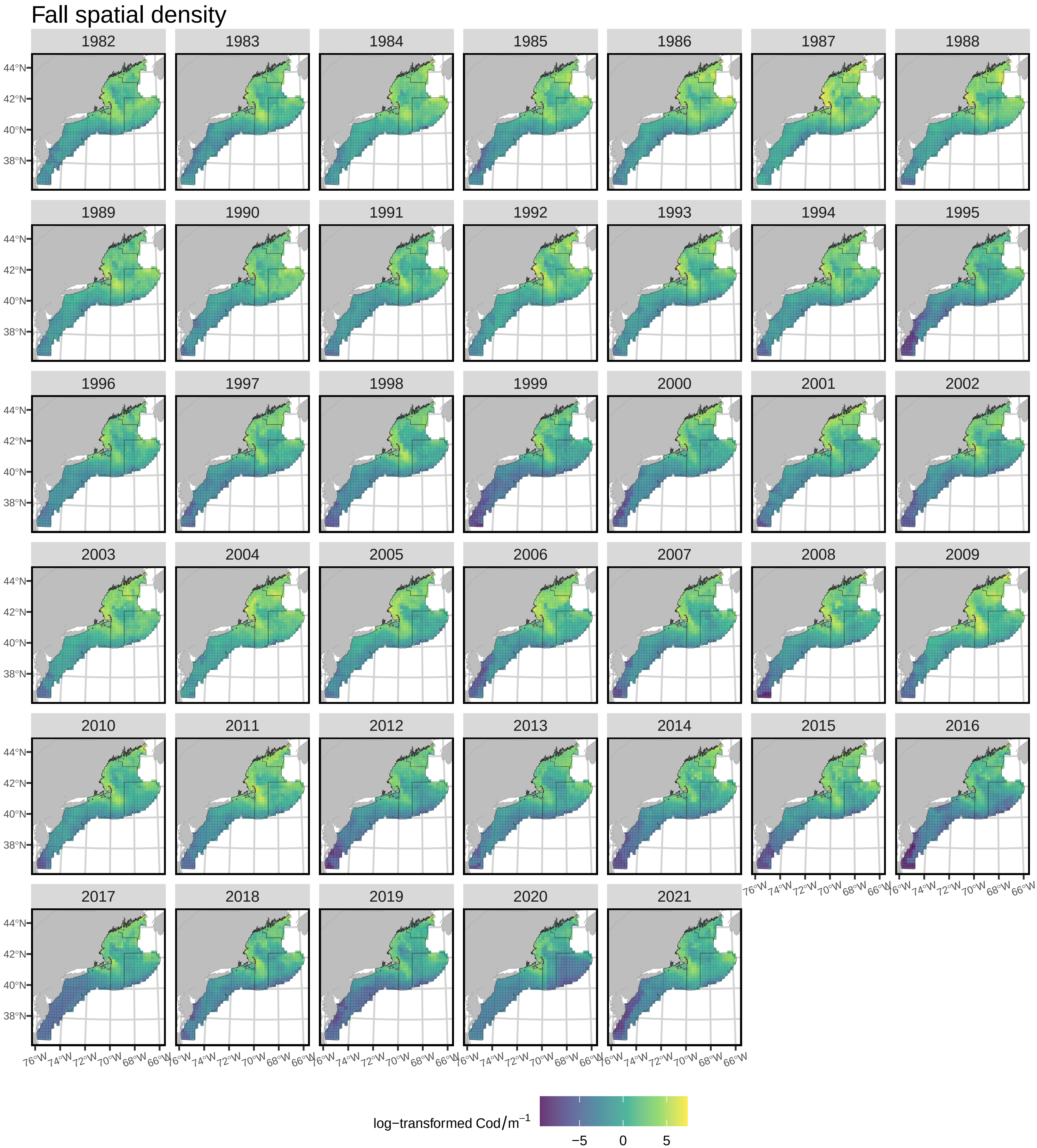

### Supplemental Figure 7a

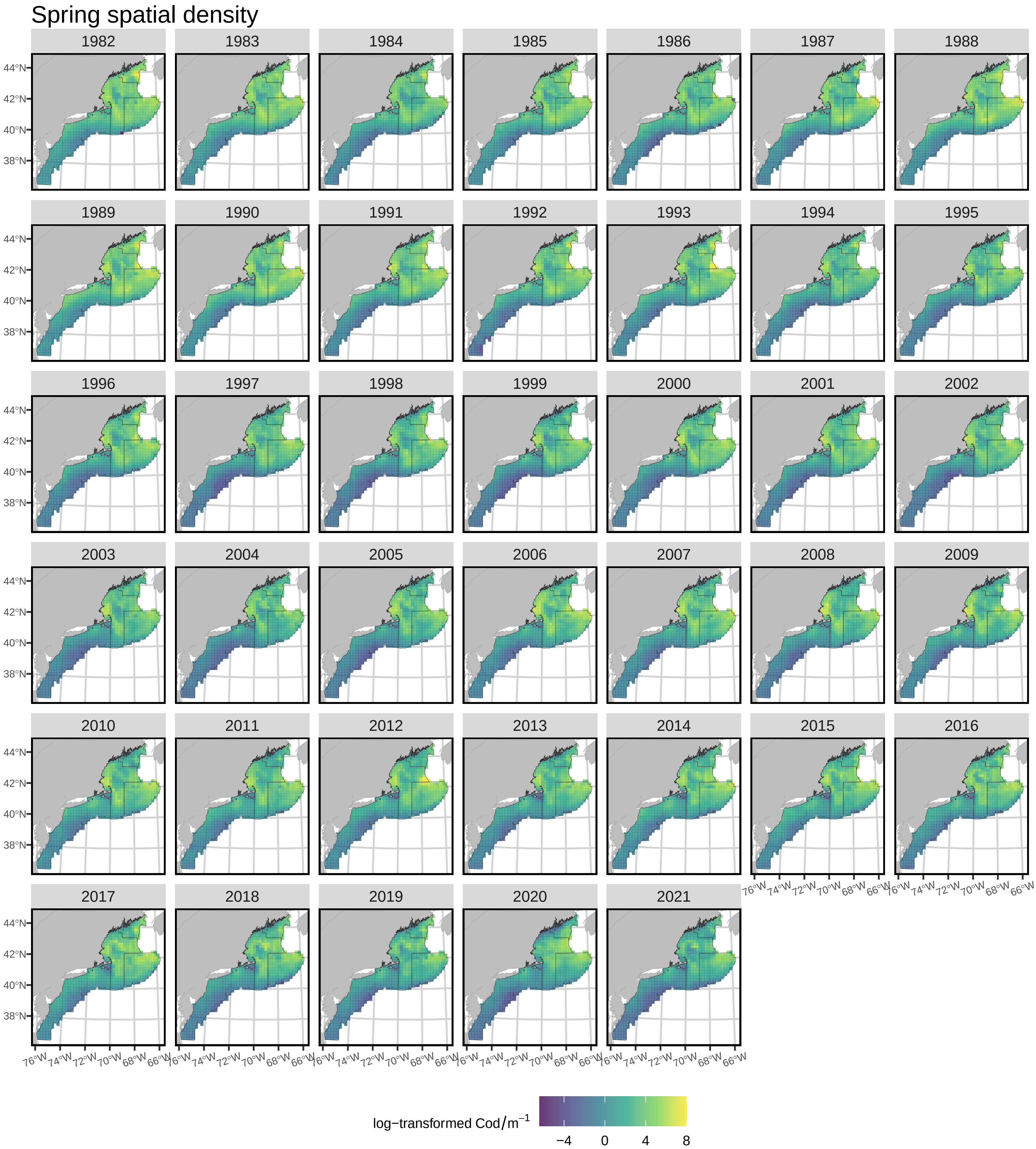

### Supplemental Figure 7b

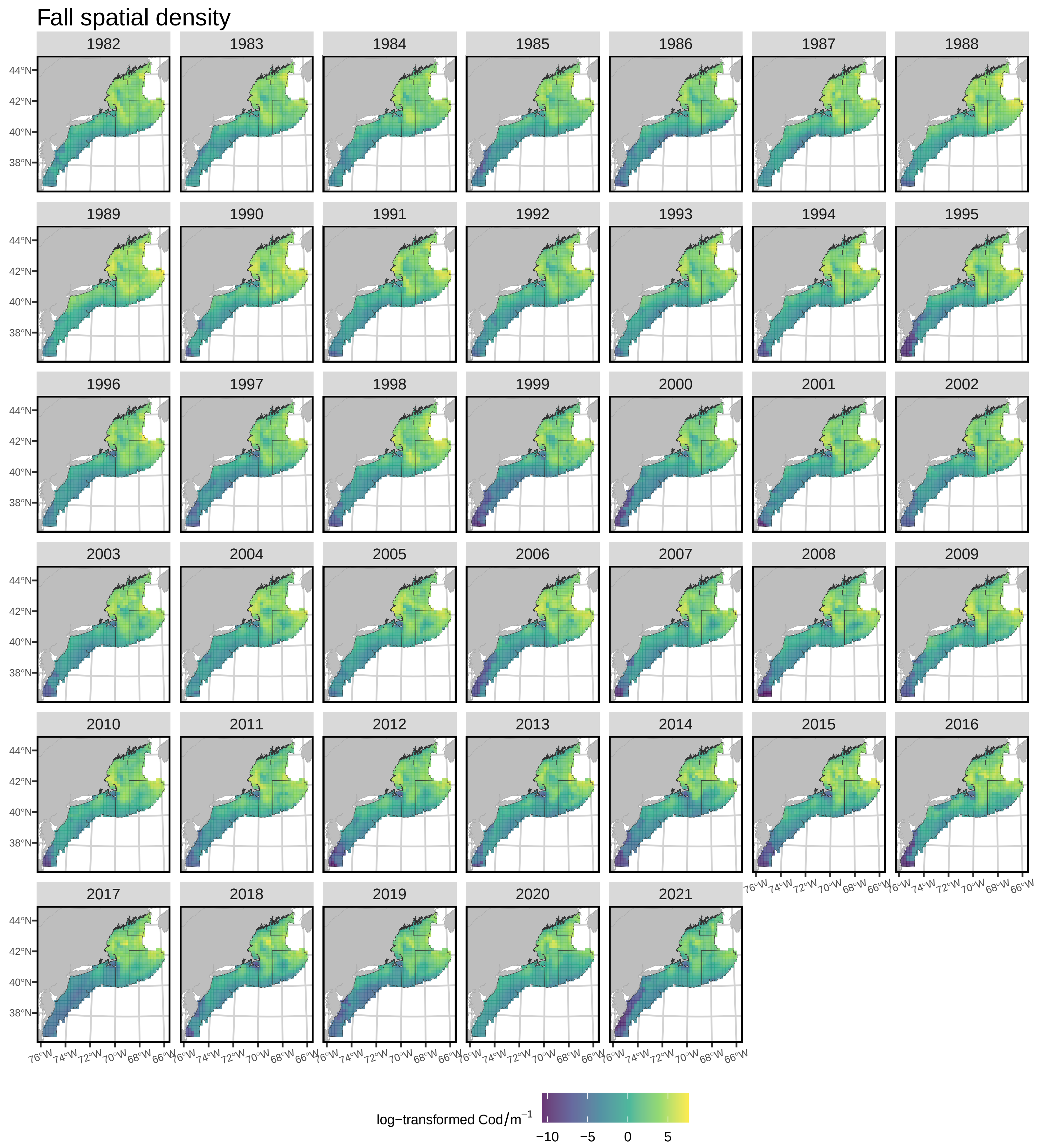

### Supplemental Figure 8a

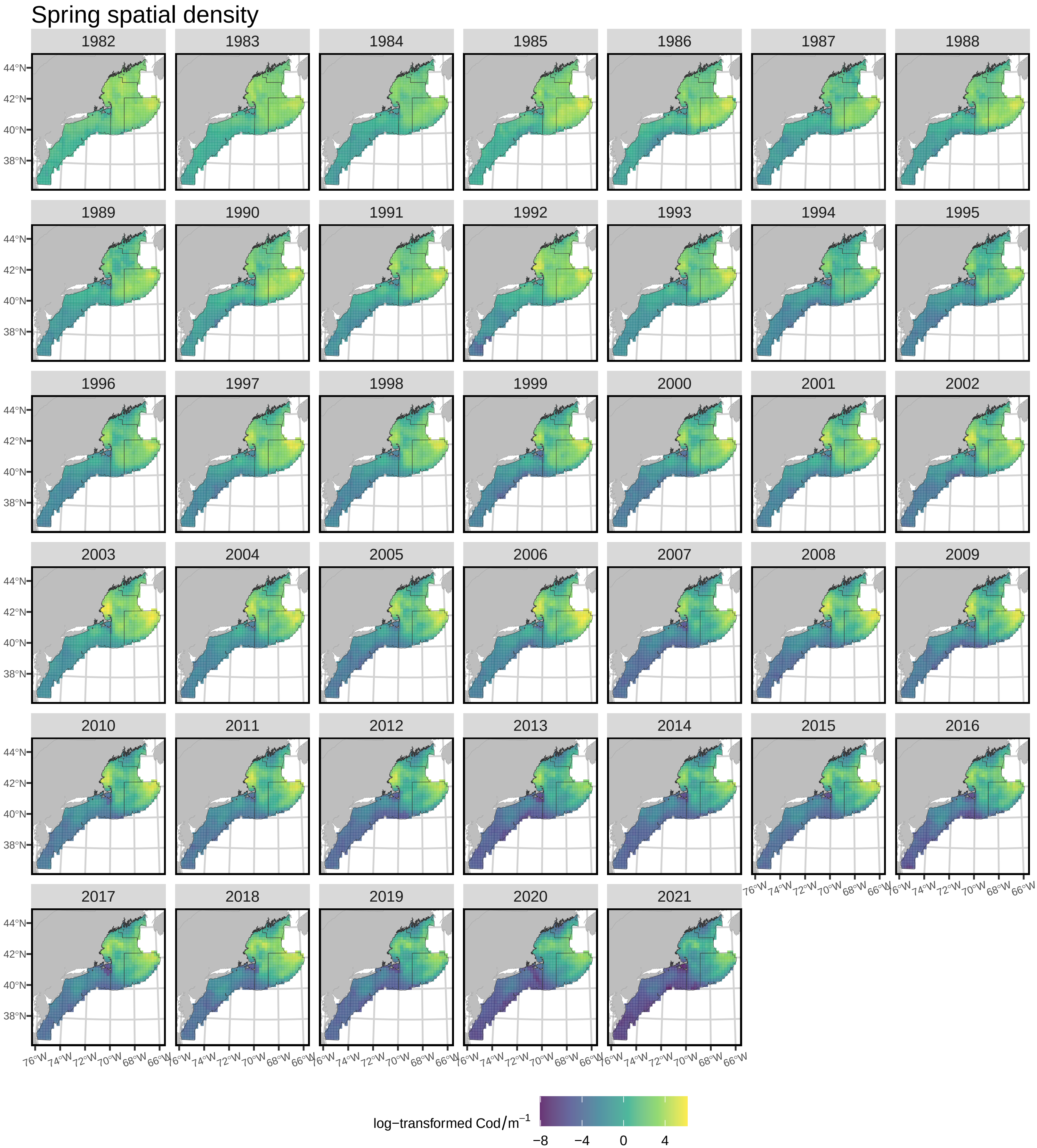

### Supplemental Figure 8b

#### Fall spatial density

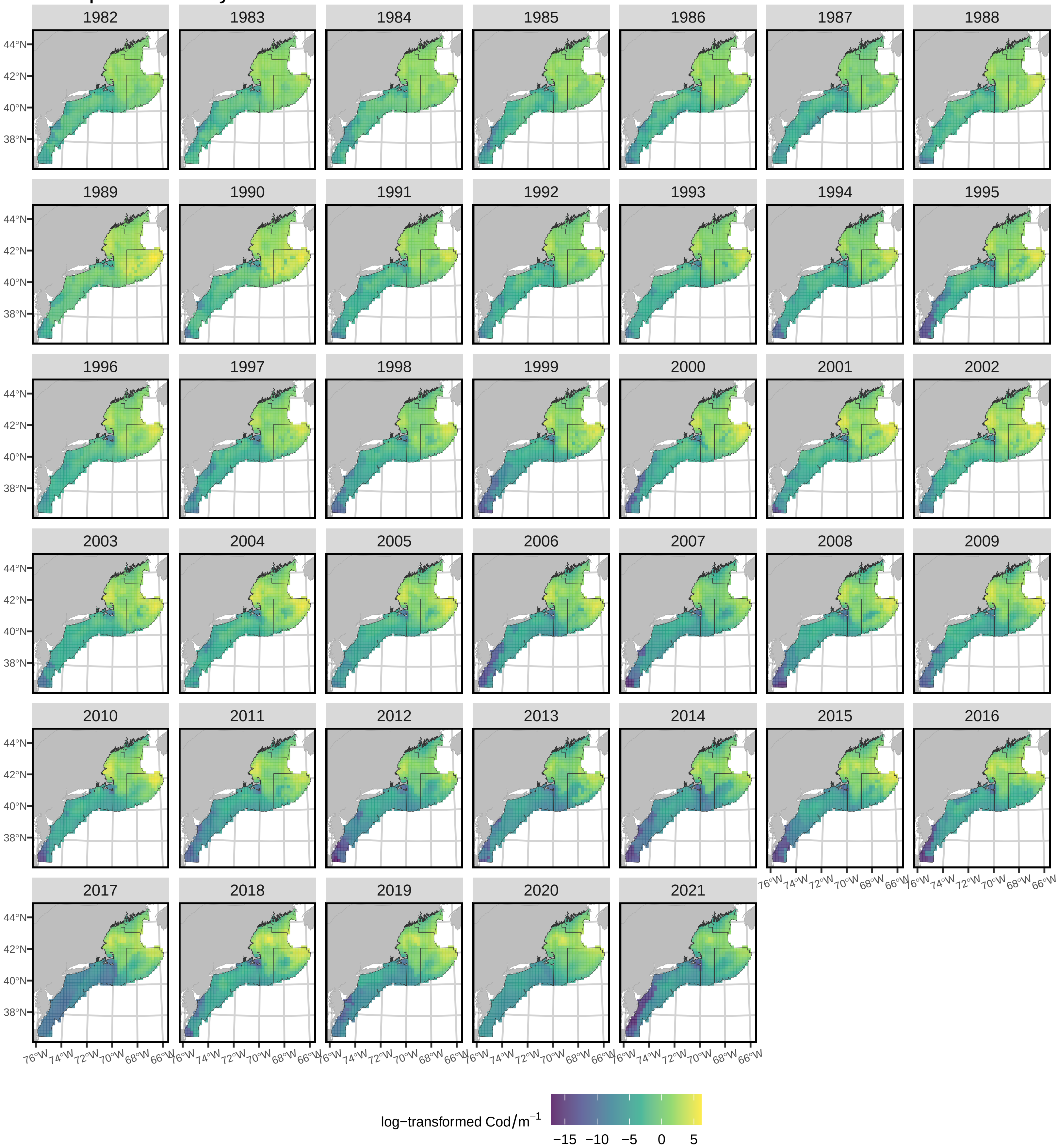

### Supplemental Figure 9

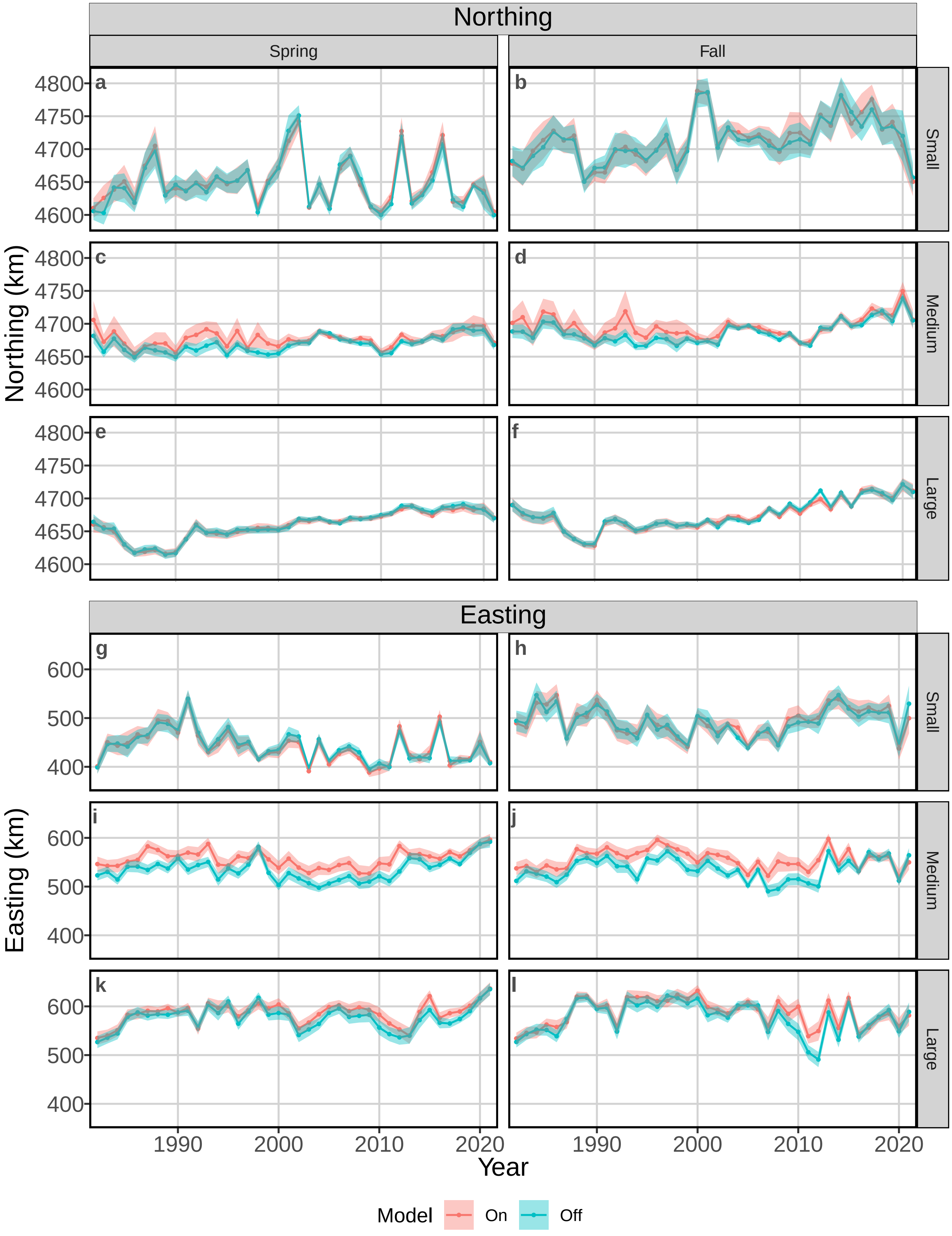
